# Left-Right Brain-Wide Asymmetry of Neuroanatomy in the Mouse Brain

**DOI:** 10.1101/2024.06.25.600709

**Authors:** Andrew Silberfeld, James M. Roe, Jacob Ellegood, Jason P. Lerch, Lily Qiu, Yongsoo Kim, Jong Gwan Lee, William D. Hopkins, Joanes Grandjean, Yangming Ou, Olivier Pourquié

## Abstract

Left-right asymmetry of the human brain is widespread through its anatomy and function. However, limited microscopic understanding of it exists, particularly for anatomical asymmetry where there are few well-established animal models. In humans, most brain regions show subtle, population-average regional asymmetries in thickness or surface area, alongside a macro-scale twisting called the cerebral petalia in which the right hemisphere protrudes anteriorly past the left. Here, we ask whether neuroanatomical asymmetries can be observed in mice, leveraging 6 mouse neuroimaging cohorts from 5 different research groups (∼3,500 animals). We found an anterior-posterior pattern of volume asymmetry with anterior regions larger on the right and posterior regions larger on the left. This pattern appears driven by similar trends in surface area and positional asymmetries, with the results together indicating a small brain-wide twisting pattern, similar to the human cerebral petalia. Furthermore, the results show no apparent relationship to known functional asymmetries in mice, emphasizing the complexity of the structure-function relationship in brain asymmetry. By establishing a signature of anatomical brain asymmetry in mice, we aim to provide a foundation for future studies to probe the mechanistic underpinnings of brain asymmetry seen in humans – a feature of the brain with extremely limited understanding.

**Significance Statement:** The human brain shows significant left-right anatomical asymmetry. Understanding its microscopic basis has implications for studies of autism and schizophrenia, evolution, embryonic brain development, and the relationship between structure and function in the brain. One of the biggest challenges to understanding this aspect of the brain is that animal models are limited. Here we show a brain-wide twisting pattern of asymmetry in the mouse brain using over 3,500 animals from six independent cohorts. These findings provide a basis for using mice to interrogate the microscopic underpinnings of anatomical asymmetry in humans and a roadmap for exploring anatomical asymmetry in additional species.

## Introduction

A central feature of the vertebrate body plan is its bilaterally symmetrical left-right organization. Some structures, such as the visceral organs break this symmetry as a result of asymmetrical signaling during development^1^. The human brain, while generally symmetrical between sides, exhibits distinct asymmetries in both anatomy and function. Numerous cognitive abilities such as language and face recognition depend on circuitry that reliably exhibits greater activation on one side of the brain than the other, and does so in a consistent direction across the population^2–8^. Anatomically, most regions of the human brain show small population-average asymmetries of cortical thickness (CT) or surface area (SA)^9–12^, alongside a macro-scale “twisting” of the brain called the *cerebral petalia* in which the right hemisphere is shifted anteriorly relative to the left^13–17^. Anatomical asymmetry of the posterior temporal lobe (the *planum temporale*) is thought to relate to functional asymmetry of human language, although studies show this relationship between anatomical and functional asymmetries to be complex at the individual level^18–22^.

Extensive research in humans on anatomical asymmetry has identified highly reproducible patterns of CT and SA asymmetries^9,23^ as well as an anterior-posterior pattern of CT asymmetry^9,23–25^. Neurodevelopmental and cytoskeletal genes have been associated with modified anatomical asymmetry^26^, with some of these genes also implicated in a recent genome-wide association study of handedness^27^. However, fundamental aspects of human anatomical brain asymmetry remain poorly understood and challenging to address in human subjects. For example, the cellular basis of anatomical asymmetries seen in humans is still unknown, as is the mechanism behind the small shifts observed in asymmetry for individuals with autism spectrum disorder^28,29^ and schizophrenia^30,31^. Furthermore, the developmental mechanisms driving region-specific brain asymmetries remain unclear, as does the extent to which anatomical asymmetries are even present in other mammalian species.

Understanding of these questions would benefit substantially from animal models for conducting cellular-level studies of anatomical asymmetry. However, existing models have limitations which largely hinder their applicability to asymmetry in humans. In zebrafish and *Drosophila*, well-defined asymmetries respectively exist in the habenula^32–34^ and brainstem^35,36^, but the rest of the brain in these animals shows no other apparent morphological asymmetries. This raises an important question about whether these isolated examples seen in distant species are fundamentally distinct from the extensive, subtle pattern of asymmetries seen brain-wide in humans. In chimpanzees and macaques, asymmetry has been generally reported across many brain regions, but studies show limited reproducibility^37–43^. This is further complicated by the difficulty of comparing studies using different metrics (volume^37,38^, SA^25^, CT^25,44^), and different levels of granularity (voxel-based^43,44^ vs atlas-based^25,42^ results). One exception to this reproducibility challenge is the *planum temporale*, which is widely seen to be expanded in the left hemisphere in chimpanzees^25,37,45–48^ and baboons^49–51^ – similar to humans^52,53^. Yet, this represents the only well-defined example of asymmetry within those species. As a result, it has been difficult to establish a consensus pattern of asymmetry for the whole brain in any species outside of humans.

Alternatively, mice, which are a commonly used species for understanding the human brain’s organization^54^, are an ideal model organism for studying neuroanatomical asymmetry. Despite its overall appearance of symmetry, the mouse brain possesses many well-described functional asymmetries: processing of vocalizations and pitch sweeps^55–58^, visceral stimuli^59^, spatial memory^60–63^, fear^64–67^, and handedness^68^. In addition, numerous brain-wide structural imaging datasets and sensitive tools for morphometry analysis exist^69–72^, allowing for investigation into neuroanatomical asymmetries along with replication analyses between cohorts – something that is largely absent from the existing non-human literature.

Three prior studies have investigated neuroanatomical asymmetry in mice using single cohorts, but as in primates, they reached conflicting descriptions. One study by Spring et al. 2010^73^ showed an overall larger left hemisphere volume of 2.8% while a more recent study by Elkind et al. 2023^74^ identified an asymmetry pattern varying regionally without any clear trend. A third investigation by Rivera-Olvera et al. 2024^75^ further identified regional volume asymmetries but which did not resemble either of the 2 previous studies. However, this investigation also reported an anterior-posterior pattern of asymmetry: the right side being larger in the anterior with the left side larger in the posterior. The widespread availability of distinct imaging datasets from separate groups now provides the opportunity to comprehensively investigate this question in mice while accounting for the overall generalizability of the results.

Here, we investigate neuroanatomical asymmetry in mice using multiple cohorts from different imaging modalities. We found that anatomical brain asymmetry is indeed present in the adult mouse brain but is initially obscured by strong cohort-specific patterns of asymmetry without obvious trends. By instead looking for what is consistent between these groups of animals, we identified an anterior-posterior pattern of volume asymmetry where anterior regions are larger on the right and posterior regions are larger on the left. The results together indicate a global “twisting” of the brain in mice, similar to the cerebral petalia, rather than the distinct local asymmetries observed in humans. We thus propose mice as a novel mammalian animal model for investigating neuroanatomical asymmetry at the microscopic level. Future understanding of the genetic and cellular underpinnings of this asymmetry signature may provide insight into observations seen in humans.

## Results

### Pipeline Overview

We obtained brain-wide imaging data from six different published studies performed by five different research groups (**Table S1**). These studies used mice of different genetic backgrounds, with drastically varying numbers of animals (n = 23 to 2,992). We used three high-resolution serial histology studies (Serial Two-Photon Tomography; STPT) and three magnetic resonance imaging (MRI) studies. Animals of a single study are referred to here as a “cohort”. To investigate anatomical asymmetry, we used automated image analysis to measure volume in each atlas-defined brain region (**Fig. 1A**). We used the well-established software ANTsPy^71^ to perform automated linear and non-linear image registration, which aligned each sample image to a symmetrical, three-dimensional (3D) reference template. This produces a 3D image (Jacobian determinant field) in which each pixel represents the local volumetric change between the original sample and the reference template. We then obtained region-specific left-right asymmetries by integrating over this field in combination with the pixel-wise atlas annotations (Common Coordinate Framework version 3; CCFv3).

**Figure 1.**
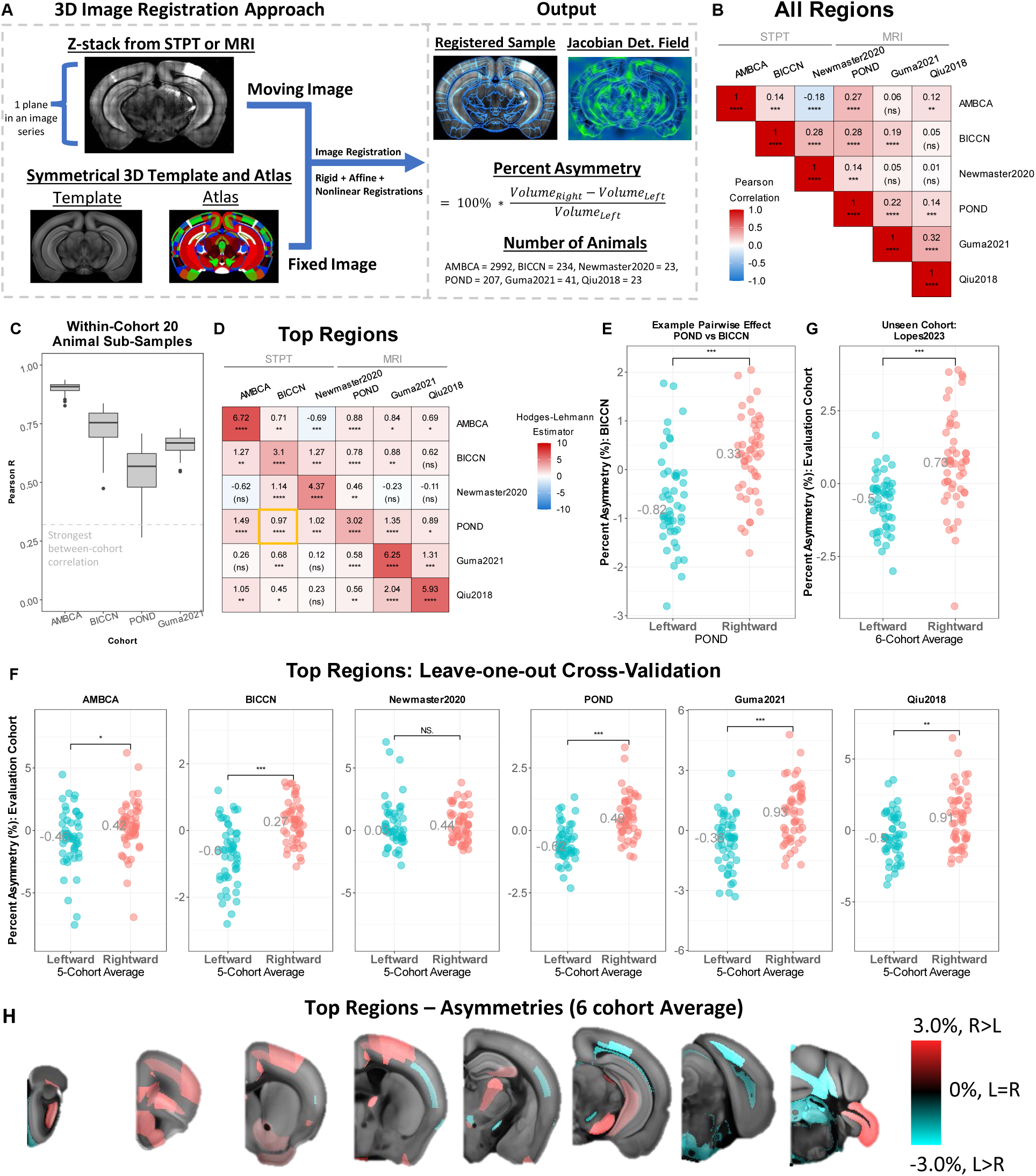
Reproducible Region-wise Asymmetry Patterns in the Adult Mouse Brain. **A)** Experimental workflow. **B)** Matrix of region-wise asymmetry comparisons between cohorts for all brain regions. Cell value is Pearson correlation between the 2 cohorts for asymmetry of all regions. Second line in cell indicates significance of the correlation. **C)** Distribution of Pearson R values from sub-sampling 20 animals and comparing the sub-sampled asymmetry pattern with either the full cohort (AMBCA,BICCN,POND), or the remaining 21 animals (Guma2021). Dashed line is largest non-diagonal value in B). **D)** Matrix of region-wise asymmetry comparisons between datasets using top 50 leftward and top 50 rightward regions. Cell value is the median of differences (Hodges-lehmann estimator). Top regions are defined in the cohort along the row and then evaluated using the cohort in the column. Gold rectangle is example shown in E). **E)** Example of top regions categorical scatter plot. Each point is 1 brain region. Top regions were defined using the POND dataset and evaluated using the BICCN dataset. Grey value indicates the median asymmetry in the evaluation cohort for that set of brain regions. **F)** Leave-one-out analysis where title of plot indicates which cohort was left-out and used as the evaluation cohort. Top regions were defined using each 5-cohort average. **G)** Evaluation of 6-cohort average in previously un-tested Lopes2023 cohort (n = 48 animals). **H)** Montage of top 100 regional asymmetries in the 6-cohort average pattern. **D-G)** Statistics are Wilcoxon rank-sum test. * is p < 0.05; ** is p < 0.01; *** is p < 0.001; **** is p < 0.0001.

### Reproducible Volume Asymmetries Exist Across Independent Cohorts

Our investigation of asymmetry initially considered each cohort individually. Doing so revealed a large number of significantly asymmetric brain regions in each of the 4 largest cohorts as well as a substantial enrichment for significant asymmetries in the 2 smallest (**Fig. S1**). However, when we attempted to correlate these region-wise patterns across cohorts, we found only weak consistency between them (**Fig. 1B)**. In contrast, sub-sampling 20 animals from a given cohort showed much stronger consistency within that cohort (**Fig. 1C**). We then concentrated on the 50 most significant leftward (left > right) and 50 most significant rightward (right > left) asymmetric regions from each cohort, out of 615 atlas regions, asking whether the most significant asymmetries from one cohort could predict the direction of asymmetry for those same regions in a second cohort. We observed such a result in 73% of cross-cohort comparisons (**Fig. 1D-E**), demonstrating a degree of predictive capability but which was generally noisy. Overall, these asymmetry patterns, despite their significance, indicate strong cohort-specific effects, with little linearity in the relationship across cohorts.

To instead identify more consistent asymmetries, we reasoned that the average asymmetry pattern of several cohorts together is likely to provide a more reproducible set of results than any one cohort individually. We thus performed a leave-one-out cross-validation, averaging the summary patterns of asymmetry from five cohorts and comparing the resulting values to the remaining sixth cohort, repeating this procedure for each combination of five training cohorts and 1 testing cohort. In five of the six possible combinations, the top 50 leftward and top 50 rightward asymmetries, defined using the five-cohort average, significantly predicted the direction of asymmetry in the remaining sixth cohort (**Fig. 1F**). The last combination showed a non-significant trend in the expected direction (likely non-significant due to the high rate of tissue defects present in the images from that study, combined with its low number of animals). This result means that on five of six unique occasions, the top asymmetries obtained through a five-cohort average generalized well to the remaining sixth cohort. By then taking the average asymmetry of all six cohort summaries together, we obtained a final, high-confidence description of asymmetries which are generalizable across independent cohorts.

We lastly confirmed that this result extended to new animals by identifying a seventh, previously un-analyzed cohort of mice from additional public data. We found that our final six-cohort average successfully and significantly predicted the pattern of asymmetry in this seventh previously un-used cohort (**Fig. 1G**). We further found that this result could be obtained with 80% power when using only 16-animal sub-samples from this cohort, indicating that for future studies, smaller-sized datasets can be used (**Fig. S2**). Overall, the magnitude of the most significant asymmetries in the six-cohort average ranges from 0.3% to 2.8% difference in regional volume between sides (**Fig. 1H, Dataset S1**). The biological implications of this asymmetry pattern are explored in **Fig. 4-7**, after the control experiments presented next (**Fig. 2-3**).

### Brain Asymmetry in Mice Is Not Due to Technical Artifacts or Biases

We first sought to validate that our morphometry analysis can recapitulate previously reported results in a non-asymmetry context. We therefore used the same volume data generated above to calculate regional male-female differences in volume (**Fig. 2A**). Previous studies in mice have defined a clear signature of male-female neuroanatomical differences – a larger bed nuclei stria terminalis, medial amygdala, and medial preoptic area in males^74,76,77^ – which we utilized as a positive control analysis. We found that male-female differences are highly reproducible across all 6 original cohorts, and we identified this expected regional signature as the top result (**Fig. 2B-G**). Additionally, we quantitatively compared our data to previous results by Elkind et al. 2023^74^ who used one of the same cohorts as us (AMBCA). For this cohort, our results were consistent with theirs, both for sex differences as well as for asymmetry (**Fig. 2H-I, Fig. S3**), despite our groups using independent pipelines. Overall, these results confirm that we are recapitulating established findings.

**Figure 2.**
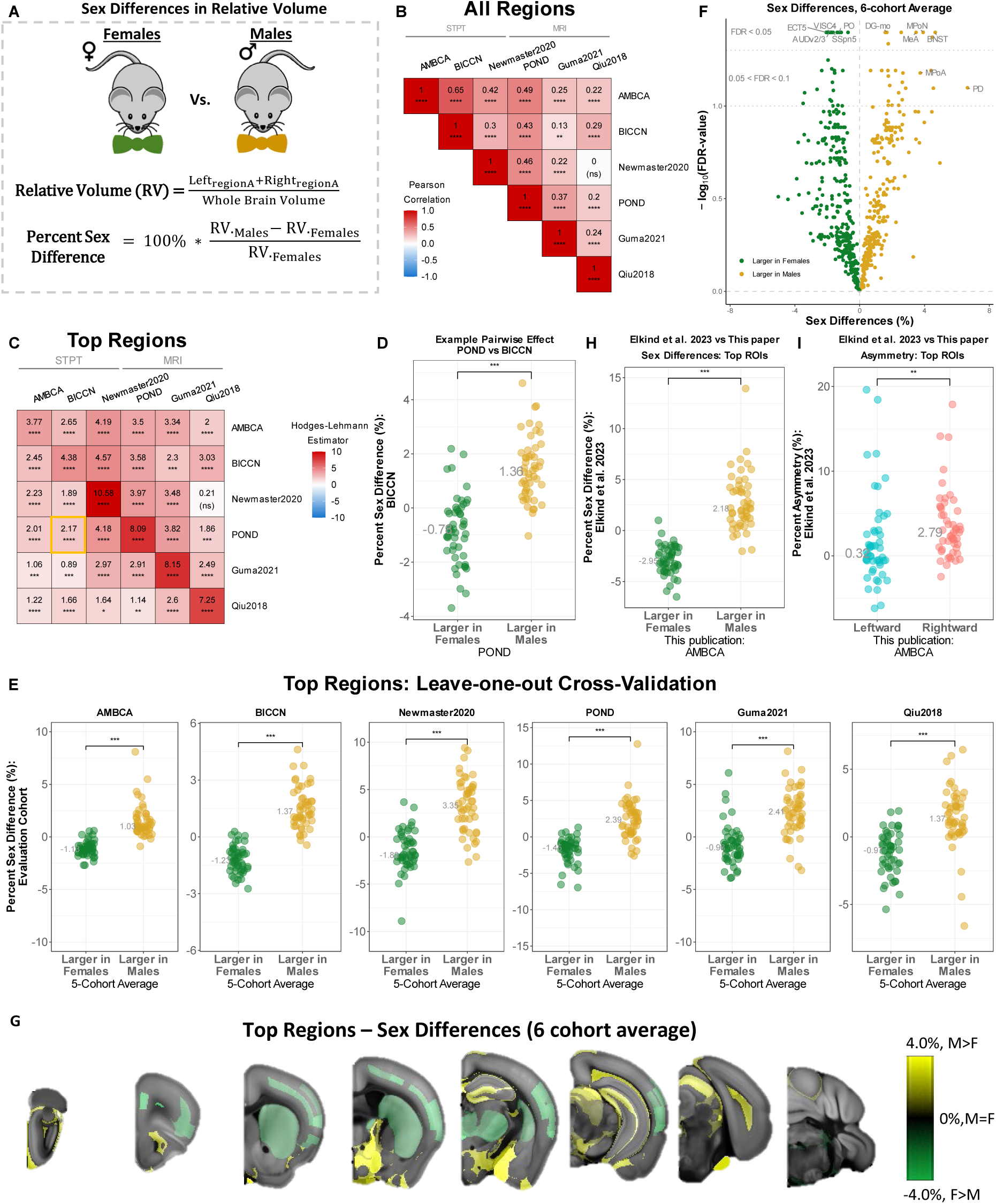
Same Jacobian Maps Reveal Highly Reproducible and Established Sex Differences. **A)** Experimental overview. **B)** Matrix of region-wise sex-difference comparisons between cohorts for all regions. **C)** Matrix of region-wise sex-difference comparisons between cohorts using top 50 male-larger and top 50 female-larger regions. Value is median of differences (Hodges-lehmann estimator). Top regions are defined from the cohort along the row and are then evaluated using the cohort in the column. Gold rectangle is example in D. **D)** Example of top regions categorical scatter plot highlighted by gold rectangle in C). Each point is 1 brain region. Top regions were defined using the POND dataset and evaluated using the BICCN dataset. Values indicates the median sex difference in the evaluation cohort for that set of brain regions. **E)** Leave-one-out analysis where title of plot indicates which cohort was left-out and used as the evaluation cohort. Top regions were defined using each 5-cohort average. **F)** Volcano plot of significant sex differences using 6-cohort average. **G)** Montage of top regional sex differences in the 6-cohort average pattern. **H-I)** Comparison of this publication’s AMBCA **H)** sex differences and **I)** asymmetries to results of Elkind et al. 2023, who used the AMBCA as well. **C-E, H-I)** Statistics are Wilcoxon rank-sum test. * is p < 0.05; ** is p < 0.01; *** is p < 0.001; **** is p < 0.0001.

We next sought to assess whether any bias in the image processing pipeline is contributing to our observed asymmetries. To do this, we reversed the left-right orientation of 1 STPT cohort (BICCN) and 1 MRI cohort (POND), as well as artificially symmetrized them, and re-ran our analysis. In the STPT cohort, left-right flipping the original images prior to any processing almost entirely inverted the asymmetry pattern (**Fig. 3A**). The symmetrized images similarly showed much smaller asymmetries than non-symmetrized images, but which, interestingly, were still highly significant (**Fig. 3B-C; Fig. S4A**). Nonetheless, this data shows that asymmetries in our non-symmetrized STPT cohorts depend almost entirely on the image left-right orientation, an important result which demonstrates that there is no left-right bias in the pipeline to drive asymmetries. Similarly, in the MRI cohort, we found that asymmetries also depended strongly on the left-right orientation but included some noise (**Fig. S4A-D**), likely from an MRI-specific algorithm (the N4 intensity inhomogeneity correction step), which, when given a pair of identical but flipped input images, does not produce perfectly flipped output images (**Fig. S4E-F**). We tested whether this effect is contributing to the overall asymmetry pattern by comparing the average of mouse MRI cohorts to the average of mouse STPT cohorts (**Fig. 3D-E**) – the latter of which do not use this MRI-specific algorithm during their processing. We found that asymmetry generalizes between these two fundamentally distinct image modalities and thus asymmetry is not a product of this noise in the pipeline or any MRI-specific or STPT-specific artifact – such as *ex cranio* (STPT) vs *in cranio* (MRI) imaging. Overall, the results indicate that our observed asymmetries depend predominantly on the image left-right orientation and not on pipeline or tissue processing biases.

**Figure 3.**
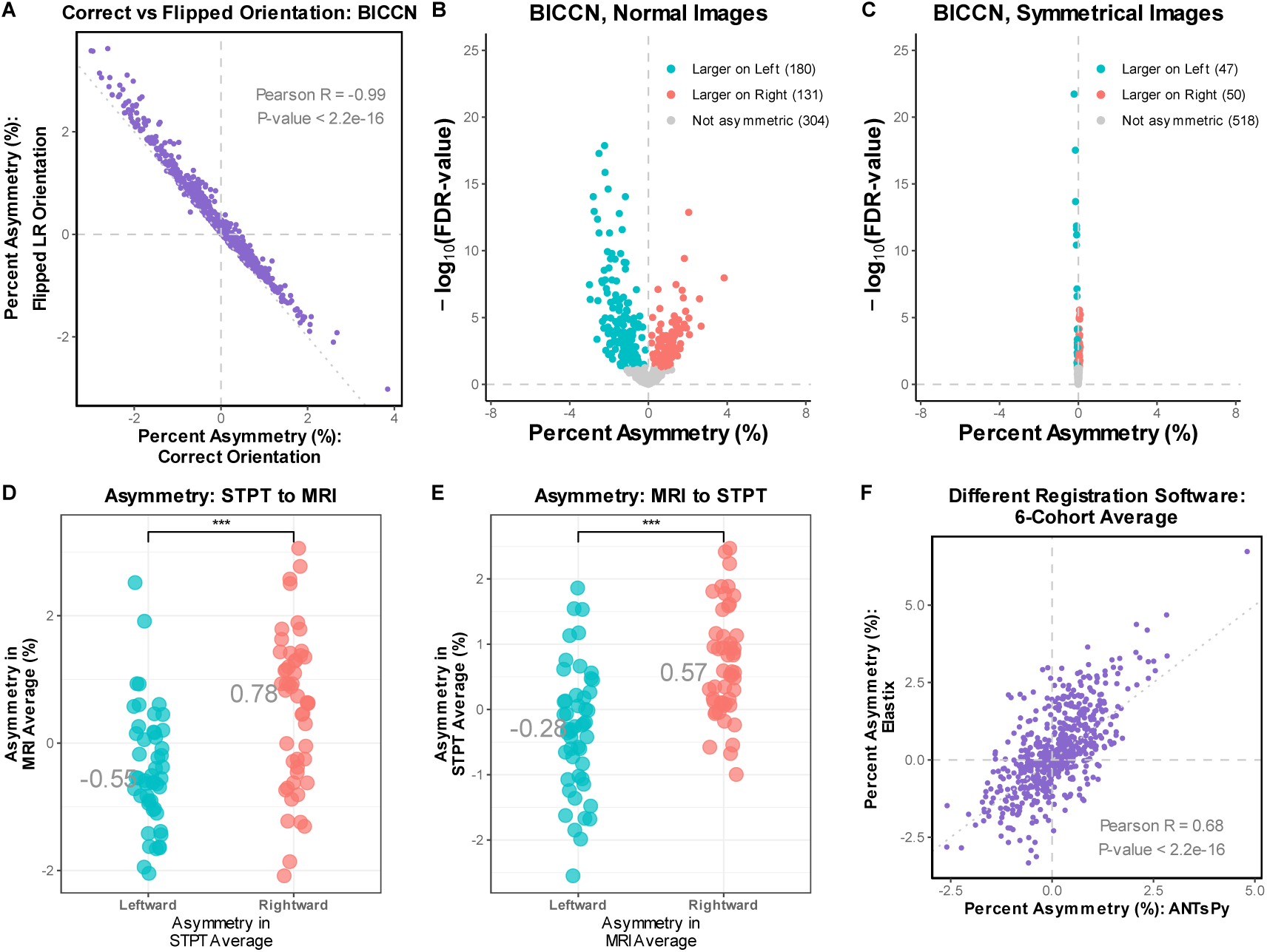
Asymmetries Are Not Due to Technical Artefacts or Biases. **A)** Comparison of BICCN region-wise asymmetry pattern between original and left-right reversed images. Each point is a brain region where the value is the mean asymmetry for that region across all animals in each analysis. **B-C)** Volcano plot of asymmetry for BICCN cohort where each point is a brain region. **B)** Analysis using normal images (same as **Fig S1B**). **C)** Analysis using synthetically symmetrized samples. **D)** Comparison of STPT-specific average asymmetry pattern (X-axis) in the MRI-specific average (Y-axis) using top regions. **E)** Comparison of MRI-specific average asymmetry pattern (X-axis) in STPT-specific average (Y-axis) using top regions. **D-E)** Statistics are Wilcoxon rank-sum test. *** is p < 0.001. **F)** Comparison of region-wise asymmetry pattern in 6-cohort average between ANTsPy and Elastix.

Finally, we assessed whether our results generalize to a different image registration software – Elastix^72^ – which uses a different transformation model from the ANTsPy registration package used above. We also used different loss functions between these software. Re-registering each sample using Elastix revealed a similar pattern of asymmetry to that observed with ANTsPy (**Fig. 3F, Fig. S5**). This result makes the prospect of an overall registration error unlikely and rules out the possibility that asymmetry is a software-specific result. Altogether, our results argue strongly in favor of a population-wide pattern of directional asymmetry existing in the adult mouse brain, robust to numerous biological and technical differences between cohorts, independent pipelines, image modalities, and registration packages.

### Top Asymmetries Are Comprised of All Major Anatomical Structures

Having established that anatomical asymmetry is indeed present in the mouse brain, we turned our attention to interpreting the regional pattern of asymmetry. Inspecting the set of top asymmetries reveals a diverse collection of regions spanning each major division of the mouse brain (**Fig. 4A**). We saw significant enrichment of atlas labels within the pons and cerebellum among the top set of asymmetries (**Fig. 4B**). However, outside of these areas, we saw no clear enrichment for any other division of the brain among our top results. Among isocortical labels, we saw no evidence of layer-specific enrichment either (**Fig. 4C**). Thus, excluding the pons and cerebellum, our set of top asymmetries mainly reflects the frequency of these structures expected by chance given their proportion in the CCFv3 atlas.

**Figure 4.**
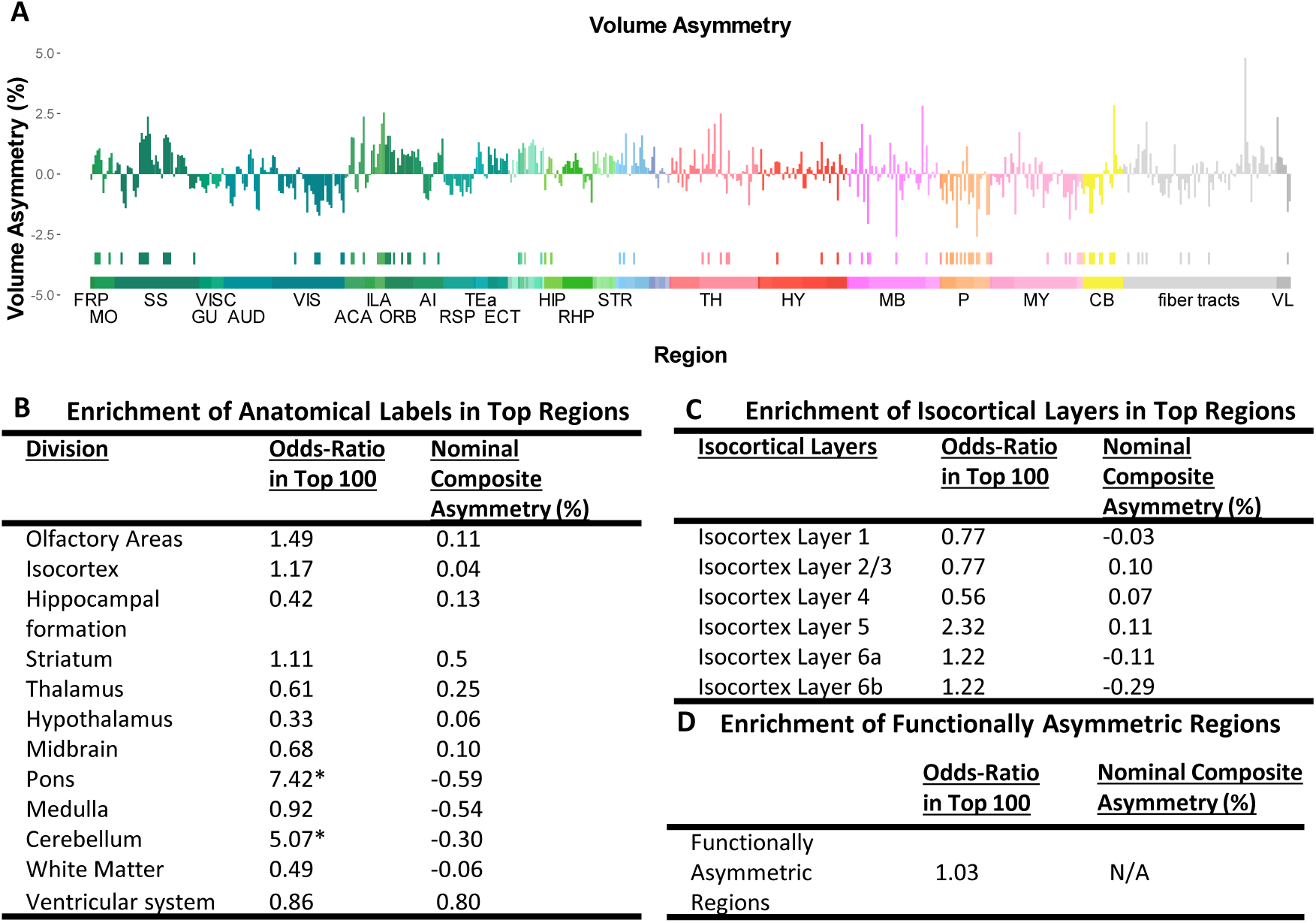
Characterization of Volume Asymmetry Pattern in the Adult Mouse Brain. **A)** Bar plot of the full Allen CCFv3 where the height of each bar along the X-axis corresponds to the nominal asymmetry in the 6-cohort average. Row of sparse ticks at approximately Y = −3 indicates whether the structure is present in set of top regions derived from the 6-cohort average: presence of tick mark indicates presence of region in top asymmetries, absence of tick mark indicates absence from top asymmetries. Color bar at approximately Y = −4.5 illustrates the broader division of tissue to which that region belongs. **B-D)** Statistical enrichment of regions within **B)** various broad anatomical divisions, **C)** Isocortical layers, or **D)** regions with established functional asymmetries in the set of the top 100 asymmetric brain regions. Nominal asymmetry of the composite structure is included as well. * indicates a significant enrichment of pons and cerebellum labels in the set of top regions at FDR < 0.05. All others are non-significant. N/A indicates not applicable.

### No Association of Anatomical Asymmetries with Functional Asymmetries

Among the top asymmetric regions, the most asymmetric structure is the Paraflocculus, with a 2.8% average rightward asymmetry (**Dataset S1**). The locus coeruleus and medial pretectal area further showed a 2.6% leftward asymmetry. Finding these regions as the top asymmetric structures was unexpected given that no evidence for lateralized function exists at these areas. Interestingly, no auditory cortical regions were found in the set of top asymmetric structures despite robust evidence of functionally asymmetric auditory processing in mice^55–58^. We therefore investigated, using the statistical enrichment approach above, whether our set of top anatomical asymmetries were indeed distinct from known functional asymmetries in mice (**Fig. S6A-B, Table S2**). We found that areas of the mouse brain with strong evidence for asymmetric function showed no association with the top anatomical asymmetries identified here (**Fig. 4D, Fig. S6C, Table S2**). We further saw through leave-one-out cross-validation that these functionally asymmetric regions don’t appear to drive the main effects seen in **Fig. 1F**, as they were less consistent across cohorts than the overall set of top regions (consistent only for 3 of 6 cohorts; **Fig. S6C**). Overall, these analyses show that gross anatomical asymmetries seen here are generally distinct from known functional asymmetries in mice.

### An Anterior-Posterior Pattern of Volume Asymmetry is Driven by SA but Not CT

Although we did not identify an association with known functional asymmetries, we did observe an anterior-posterior (AP) pattern of asymmetry in which anterior regions are larger on the right and posterior regions are larger on the left (**Fig. 1H)**. We quantified this by calculating the 3D centroid of each region from the CCFv3 atlas and plotting the AP component of that centroid against its asymmetry. We observe this pattern both when using the set of top regions or when using all regions in our 6-cohort average (**Fig. S7A, Fig. 5A**). Notably, this result is not restricted to the cortex, reflecting the global nature of the effect. This AP effect is stronger in the 6-cohort average than in any one cohort individually (**Fig. S7B-C, Fig. S8**), demonstrating how the AP effect is generally obscured by the cohort-specific patterns. Flipping the left-right orientation of the BICCN and POND images reverses the directionality of the pattern while symmetrizing these images eliminates it (**Fig. S7D-E**). Moreover, this effect is evident in both the STPT- and MRI-specific averages and is also present in our analysis using Elastix (**Fig. S7F-H**). We see no evidence of a similar trend for the dorsal-ventral (DV) or medial-lateral (ML) axes (**Fig. 5B, Fig. S7I**).

**Figure 5.**
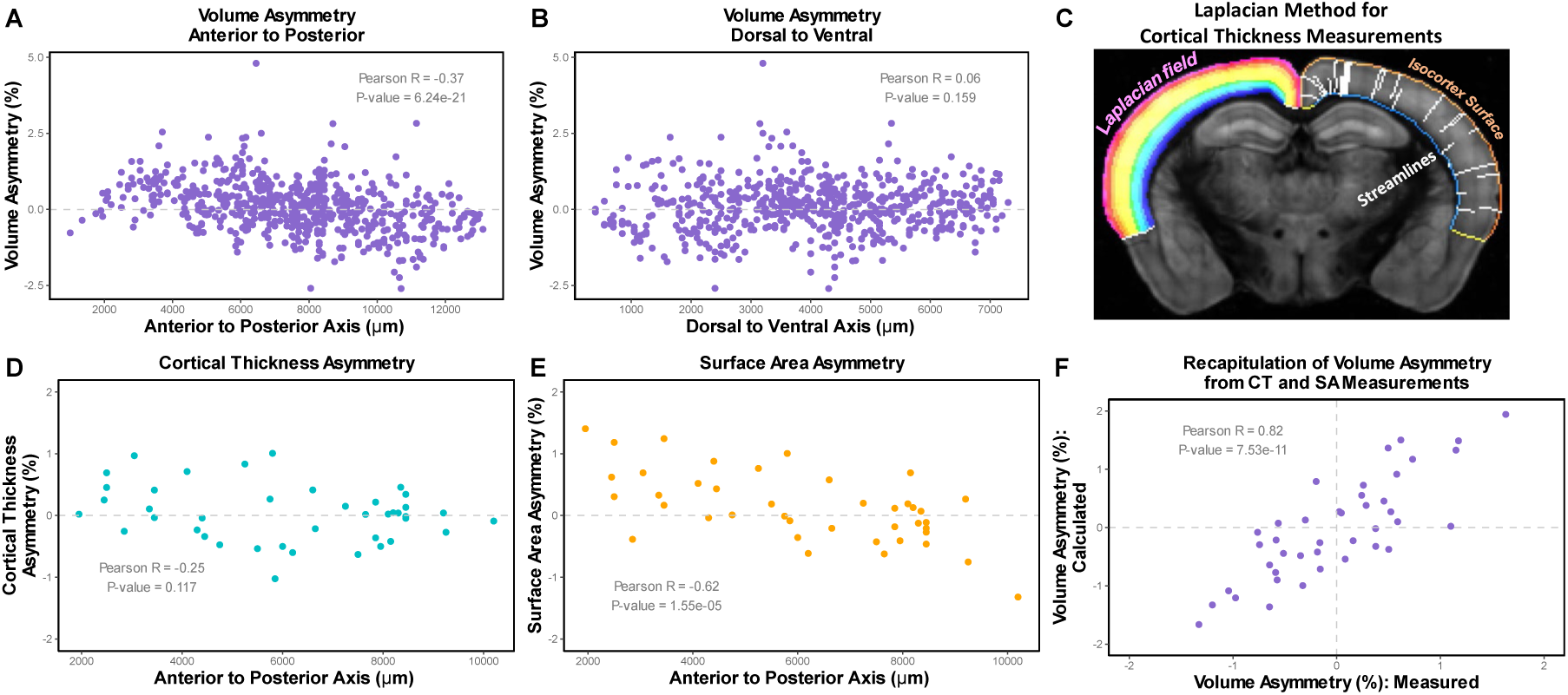
Anterior-Posterior Pattern of Volume Asymmetry is Driven by SA But Not CT. **A-B)** Average asymmetry of each brain region derived from the 6-cohort average plotted against the **A)** anterior-posterior or **B)** dorsal-ventral position of that region in the atlas. **C)** Illustration of Laplacian Method for calculating cortical thickness, during which surface area can also be collected from the “Isocortex Surface”. The cortex is modeled as 3 surfaces: the pial surface (“isocortex surface”) or the white matter-grey matter interface (blue line overlaying corpus callosum), and a special cortical geometry surface (yellow line by piriform cortex). A smooth Laplacian field is generated between the pial surface and white matter-grey matter interface. The steepest descent path through the field (streamlines) is generated from each surface pixel to the white matter-grey matter interface. **D)** Average cortical thickness per cortical territory from 6-cohort average (all layers collapsed) calculated using method in C). **E)** Average pial surface area per cortical territory from 6-cohort average. **F)** Comparison of 6-cohort average volume asymmetry per cortical territory (all layers collapsed) measured using original Jacobian-derived volume analysis with volume asymmetry calculated through separate cortical thickness and surface area measurements. In all graphs, each point is 1 brain region and the value is the mean asymmetry for the appropriate metric calculated in the 6-cohort average. All statistics are the significance of the Pearson correlation.

In humans, a CT asymmetry trend exists where anterior regions are thicker on the left and posterior regions are thicker on the right^9,23–25^. To directly assess whether our observations in mice match this reported trend in humans, we measured CT asymmetry directly in all samples. We did this by propagating the CCFv3 cortical labels onto each sample through the reverse registration and applying the Laplacian method^78,79^ (**Fig. 5C**; see supplemental methods). Although our AP volume asymmetry pattern is brain-wide, we focused on the cortex here because measurements of thickness are well-defined and because doing so best enables a direct comparison to observations in humans. We found no evidence of a pattern in mice for CT asymmetry with respect to the AP axis (**Fig. 5D**), but we also measured SA asymmetry in parallel and found a highly significant AP gradient in the same direction as our volume asymmetry pattern (**Fig. 5E**). Interestingly this gradient was in the opposite direction of the human CT pattern.

We validated these separate CT and SA measurements by using the hemisphere-specific CT and SA to re-derive volume asymmetry for each cortical territory (layers 1-6b combined into a single label per cortical area). This re-derived volume asymmetry was strongly correlated with our original volume asymmetries measured from the forward registration (**Fig. 5F**), confirming the accuracy of our measurements. Altogether, this data indicates that an AP volume asymmetry pattern exists and is driven by SA asymmetries, but not CT.

### Positional Asymmetries but No Cerebral Petalia Exists in Mice

Given the AP pattern of asymmetry observed in mice, we wondered whether this indicated a cerebral petalia-like effect. In humans, the right hemisphere protrudes ahead of the left in a brain-wide “twisting” or “warping” pattern^13–17^. Roughly two-thirds of individuals show this trend while the remaining one-third show a reversal or near-symmetry^17^. To examine this in mice, we applied an automated approach based on recent large-scale papers measuring the petalia in humans and chimpanzees^13,14^. For reference and validation, we also analyzed 4 cohorts of human subjects (positive controls) and 6 cohorts of non-human primates (NHPs; negative controls) consisting of chimpanzees, macaques, and marmosets (2 cohorts each; **Table S3**). We note that while the presence of a significant petalia in NHPs is controversial, studies agree that such an effect, if present, is much smaller in magnitude and of a weaker frequency in that species’ population than in humans^13,17,80,81^.

In humans, we found a significant right-frontal protrusion of 0.24% and a significant left-occipital protrusion of 0.70%. Both are normalized by the length of the left hemisphere and are consistent with established findings^13–16^ (**Fig. 6A-B**). In NHPs, we saw a significant right-frontal protrusion as well, but which was significantly smaller than in humans by approximately 3-fold (0.07% overall; **Fig. 6A**). In contrast, we saw no significant occipital protrusion in NHPs (**Fig. 6B**). In mice, we saw no significant protrusion for either cortical pole, with average measurements similar to the NHPs (anterior = 0.06%, posterior = 0.07%; **Fig. 6A-B**). Thus, our measurements suggest that mice do not exhibit an AP protrusion of one hemisphere with respect to the other (e.g. a cerebral petalia).

**Figure 6.**
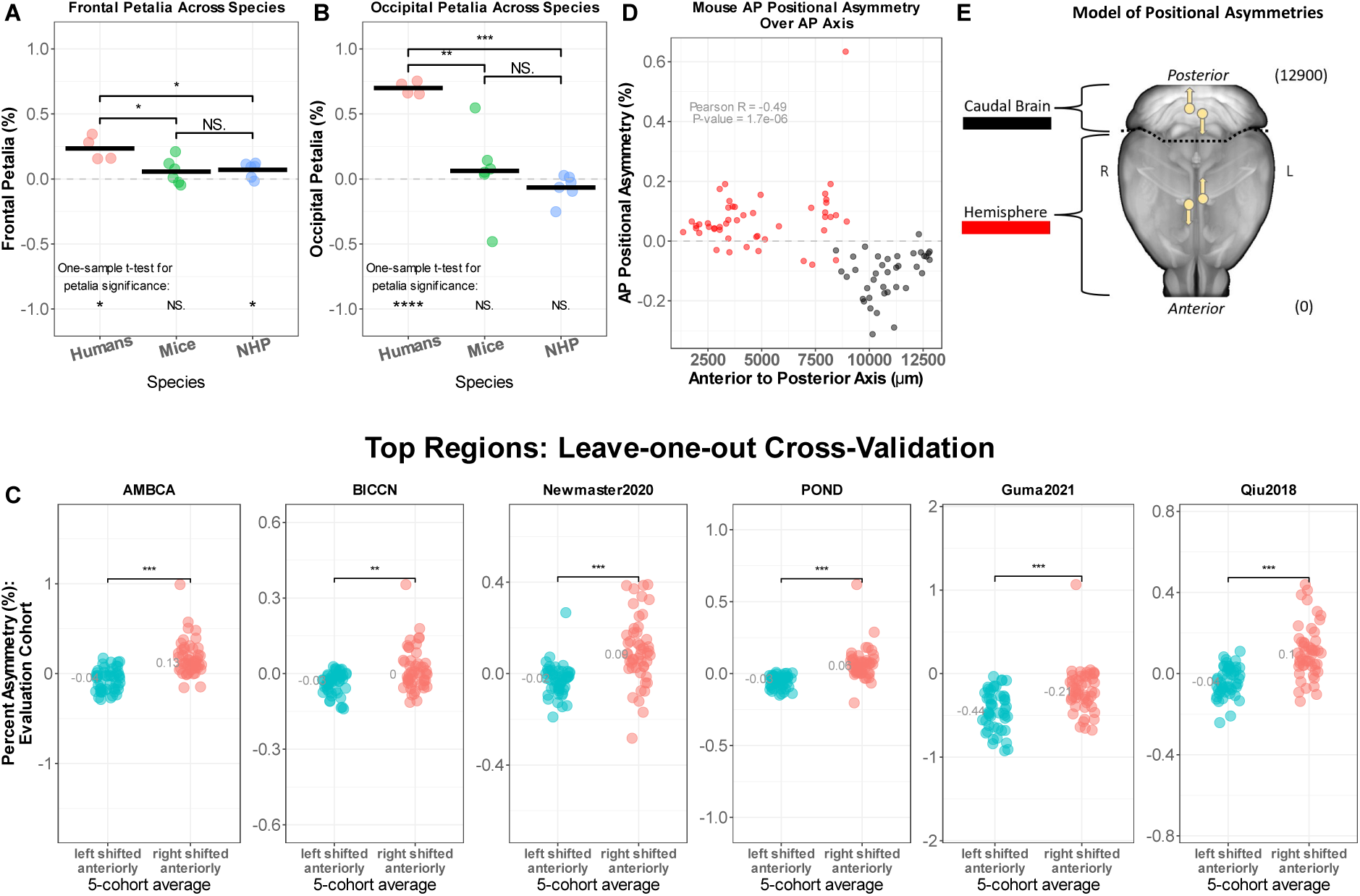
Positional Asymmetries But No Cerebral Petalia in Mice. **A-B)** Cohort-specific petalia measurements for the **A)** frontal (anterior) or **B)** occipital (posterior) poles in humans (n = 4 cohorts, ∼1400 individuals), mice (n = 6 cohorts, ∼3500 individuals), or non-human primates (NHPs; n = 6 cohorts; 2 cohorts each for macaques, chimpanzees, and marmosets; ∼1000 individuals). Each point on the graph is the average protrusion of subjects in that cohort, normalized against the length the left-hemisphere. A value of 0.2% means that the protrusion asymmetry is 0.2% the length of the left hemisphere on average. **C)** Leave-one-out cross-validation for the top positional asymmetries in all 6 cohorts. **D)** Anterior-posterior positional asymmetries derived from the mouse 6-cohort average, using the top 100 positional asymmetries. Regions that are shifted anteriorly on the right (positive values) tend to be found in the more anterior portions of the brain (“hemisphere”; red), while regions that are shifted anteriorly on the left (negative values) tend to be found in more posterior portions of the brain (“caudal brain”; black). Each point is a brain region and its color correspond to the definitions in E). **E)** Summary of positional asymmetries detected in C). “Caudal Brain” (black) is defined as midbrain, hindbrain, and cerebellum while everything anterior or dorsal to the midbrain is “hemisphere” (red). The “hemisphere” appears shifted anteriorly on the right while this trend appears opposite in the “caudal brain”. Value in parentheses are the Z-axis coordinates in pixels for anterior (0) and posterior (12900) regions. *** is p < 0.001.

Despite this, we considered whether a more sensitive approach might better reveal such an effect, leveraging the statistical power of many brain regions rather than focusing on single cortical poles. To explore this, we measured positional asymmetries across all CCFv3-annotated regions within each mouse brain, using the 3D centroids of these regions as mapped onto that animal during the petalia analysis. Cross-validation across different cohorts allowed us to assess whether the position of sets of brain regions was shifted reproducibly between sides in an AP direction. Indeed, we found highly reproducible left-right asymmetries of AP position in mice across all 6 of our cohorts (**Fig. 6C**). Regions within the cerebral hemisphere were consistently shifted anteriorly on the right side while regions within the brainstem and cerebellum were shifted anteriorly on the left side (**Fig. 6D-E**). Thus, while no significant AP protrusion asymmetry exists at the mouse cortical poles, mice do exhibit reproducible brain-wide shifts in the tissue that align with the AP volume and SA gradients.

### Evolutionary Context of Mouse Brain Volume Asymmetries

We lastly examined to what degree, if any, our observed volume asymmetries in mice were consistent with those of humans and NHPs. Initial inspection of the overall asymmetry pattern in each species revealed sharp interspecies differences (**Fig. 7A**), along with cohort-specific asymmetries for marmosets and macaques, similar to those seen in mice (**Fig. 7B, Fig. S9**). To quantitatively compare asymmetry across species, we utilized the cross-species atlas annotations from Garrin et al. 2022^82^, the marmoset and macaque cohorts used earlier, and human volume asymmetry data from Roe et al. 2023^9^. The two chimpanzee cohorts from the previous section were excluded for technical reasons (see methods).

**Figure 7.**
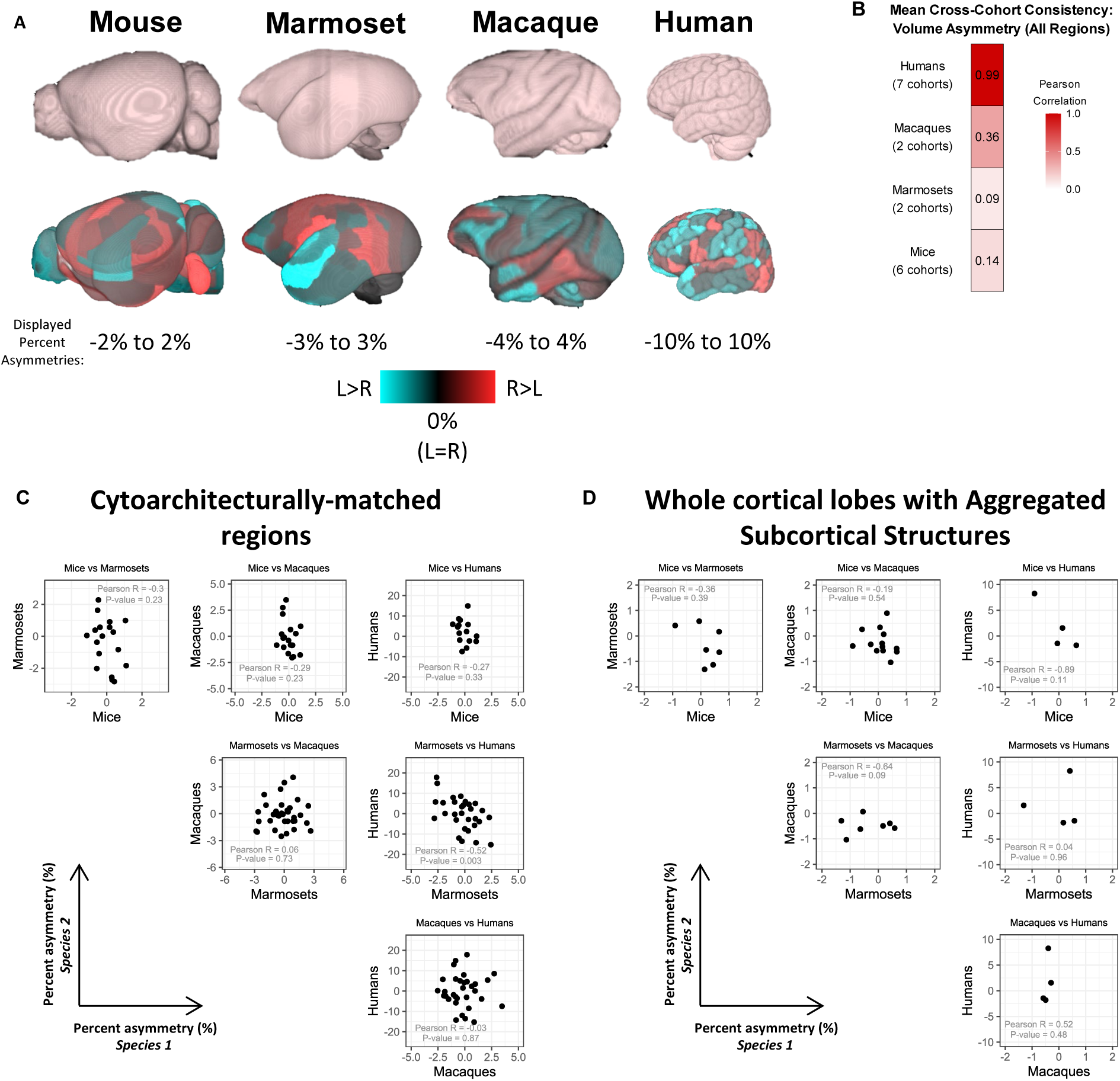
Cross-Species Comparisons of Volume Asymmetry. **A)** Top: 3D renderings of mouse, marmoset, macaque, and human brains from their respective 3D average templates. Middle: depiction of nominal volume asymmetry pattern for all regions in each atlas (regardless of significance), averaged across each cohort for that species. A minor brightness boost is added to better visualize the images. Bottom: display range of asymmetries shown for each species. **B)** The value in each cell and its color are the average cross-cohort consistency seen within a given species. Humans are the average of all 7 possible between-cohort combinations. Mice are similar but for 6 cohorts. Macaques and Marmosets have only 2 cohorts each, so the value indicates the lone Pearson correlation between the 2 cohorts in those species. **C-D)** Scatter plots of matched brain regions between species using the Garin et al. 2022 cross-species atlas. **C)** Cytoarchitecturally-matched (Brodmann) regions or **D)** summary values for whole cortical lobes and broad subcortical structures. Each species-pair has a different number of annotated regions which can be matched, hence the different number of data-points for each plot. Each point is a brain region. The value of the X- and Y-axes respectively indicate the cross-cohort average percent volume asymmetry observed in that species at that region. Human asymmetry data is only available for cortical regions.

We found weak correlations in the region-wise asymmetry pattern between any of the species analyzed, using either cytoarchitecturally matched (Brodmann) regions or whole cortical lobes with composite subcortical structures (**Fig. 7C-D**). We additionally found no evidence of an AP pattern for volume asymmetry in marmosets or macaques, but interestingly, we found an AP pattern in humans in the opposite direction of mice (**Fig. S10**) – consistent with the known CT asymmetry pattern in humans. Overall, our data indicates anatomical asymmetry to be species-specific, with little overlap between asymmetry patterns in mice, humans, and NHPs.

## Discussion

Anatomical asymmetry is a significant feature of the human brain, yet there is a lack of mammalian animal models in which to investigate its microscopic basis. Through our work, we identified an anterior-posterior pattern of volume and SA asymmetry in mice, for which anterior regions are larger and have more surface area on the right, while posterior regions are larger and have more surface area on the left. The cerebral hemisphere as a whole is shifted anteriorly on the right while the brainstem and cerebellum are shifted anteriorly on the left. Cross-cohort validation and extensive control experiments demonstrate our results, though small in magnitude, are consistent in 6 of 7 groups of animals, are free of pipeline bias, and are independent of image modality and analysis software. Our data collectively indicates that a genuine brain-wide signature of anatomical asymmetry exists in the adult mouse brain.

Considering our results in the context of humans, our findings best align with the idea of a cerebral petalia-like effect in mice, rather than the distinct local asymmetries reported in humans. Our set of 100 top asymmetries largely reflects the above anterior-posterior pattern. In this context, it makes intuitive sense that our top anatomical asymmetries lack an obvious overlap with known functionally asymmetric brain regions. The subtle asymmetries captured by our analysis appear to represent a twisting pattern manifesting across the entire brain, while region-specific asymmetries like those seen in humans or the zebrafish habenula appear to be entirely absent in mice. This apparent decoupling of anatomical and functional asymmetry in mice underscores the complexity of the relationship between anatomical and functional asymmetries reported in humans^18–22^. To-date, the basis of functional asymmetries in mice has been identified at the levels of the synapse^55,58^, protein^83^, and gene expression^55,56,59^, but our data does not support the presence of a gross anatomical substrate.

One possible explanation for the emergence of macroscale asymmetries is the molecular chirality of cytoskeletal proteins. A recent GWAS study identified enrichment for tubulin genes as genetic modifiers of anatomical asymmetry in humans^26^. In *Drosophila*, the actin cytoskeleton shows an intrinsic handedness that is sufficient to polarize a cell’s planarity and yield whole-organ and whole-organism helical-twisting^84^. Neurite growth cones in rodents have been similarly observed *in vitro* to twist in a right-screw (clock-wise) direction in a manner dependent on the actin cytoskeleton^85–87^, although whether this occurs *in vivo* is unknown. Our observations here are consistent with such a mechanism occurring in mice.

An alternative explanation could be that asymmetry emerges through a subtle timing difference in development between the left and right hemispheres. In humans, cortical folding in the temporal lobe begins 1-2 weeks sooner on the right than on the left^88–90^, and two recent studies found that transcriptome-wide changes occurring with age in the early fetal brain were slightly more advanced on one side than the other^91,92^. Anatomical asymmetries are theorized to be a “record” of this developmental timing delay^93^. Whether such an effect is occurring in mice remains unknown as does the potential mediator of such a timing delay. However, having mice as a novel animal model of anatomical brain asymmetry can be a powerful tool to investigate this and other hypotheses.

A longstanding area of interest in humans is the relationship between asymmetry of the visceral organs and asymmetry of the brain. While most evidence points toward separate mechanisms conferring asymmetry in these contexts^94–99^, one potential exception is the cerebral petalia. In humans, individuals with reversed visceral organ asymmetry (*situs inversus*) show a high frequency of a reversed cerebral petalia^95–97,100^, although the number of subjects studied in this context remains low (45 total over four studies). This emerging idea is particularly interesting in light of our observation of a similar torsion-like effect in the mouse brain. Two prior studies in mice that reversed visceral organ asymmetry observed the loss of a functional asymmetry in the hippocampus^62,101^. Thus, we wonder whether the signature of neuroanatomical asymmetry we observe here would be reversed in mice with *situs inversus*. The rarity of such individuals in humans makes this relationship challenging to interrogate (hence the low number of subjects thus far), but research using our novel signature of neuroanatomical asymmetry in mice would provide a critical datapoint for better understanding this relationship.

Toward the goal of building consensus in the mouse asymmetry literature, our study makes significant strides by drawing together the seemingly unrelated findings of Elkind et al. 2023^74^ and Rivera-Olvera et al. 2024^75^. We quantitatively replicate the volume asymmetry results of Elkind et al. while we observe the same AP asymmetry trend as Rivera-Olvera et al. Our multi-cohort analysis also clarifies the interpretation of both studies by demonstrating that apparent region-specific asymmetries in mice should be understood as part of a global pattern, rather than isolated expansions of specific structures. However, our study cannot account for the left-hemisphere bias reported in Spring et al. 2010^73^; perhaps it stems from a residual global intensity or distortion bias from the MRI. In our work, we observed a global bias in positional asymmetries of the Guma2021 cohort, with a global trend toward regions shifting anteriorly on the left, but the relative difference in asymmetry between anterior and posterior regions remained evident. Such a true global bias in the brain appears doubtful to us when considering the other cohorts in our analysis.

For future studies attempting to replicate our results, the asymmetry for the top 100 regions can serve as a direct test of replicability by asking the following question: when measured in a new set of animals, do the leftward and rightward asymmetries defined in this study vary significantly from each other in the predicted direction? (e.g. **Fig. 1F**). A significant difference in this context in the predicted direction provides evidence of replication. This approach can also be used for our reported positional asymmetries.

Our focus here on sets of asymmetries highlights the probabilistic nature of our results. The sets of leftward and rightward asymmetries we define are repeatable across independent cohorts, however, individual regions within these sets cannot be considered significantly asymmetric on their own. Nonetheless, defining sets of asymmetric regions ultimately ends up being a strength when applied to future studies: an investigation into the cellular properties of asymmetric regions is likely to be more powerful if the results are applicable over many regions – even with some added noise – rather than in only a few focused regions.

A surprising finding from our study is the significant asymmetries present in most mouse imaging cohorts but which tend to be cohort-specific. This variability contrasts with studies of sex differences, where effects in one cohort are consistently reproduced in another. Since this cross-cohort variability is specific to left-right comparisons, we believe it is unlikely to arise from factors affecting both hemispheres equally, such as image modality, number of subjects, or original image resolution. Similarly, biological influences – like age, genetic variation, the male-female ratio – appear to have only minor impacts on human brain asymmetry and would not account for the dramatic differences seen between cohorts^9,23,26,102^. Instead, we suspect technical factors, which do not impact both sides equally, may be responsible, although further research is needed to confirm this. For STPT samples, causal factors could be the intensity vignetting in the tiled images. For MRI samples, this could be residual shading biases or residual geometric distortion, among other possibilities. In principle, these technical issues could create cohort-specific local minima during the registration’s optimization process that could trap a more “aggressive” image registration – especially given the small magnitude of the asymmetries. An alternative explanation could involve husbandry-specific factors driving environment-dependent asymmetries. However, this seems unlikely given that the two marmoset cohorts we analyzed showed significant and largely distinct asymmetry patterns, despite being raised in the same environment at the same institution (personal communication, acknowledged individual J.Hata). Ultimately, our cross-cohort approach proved advantageous to a single-cohort analysis, allowing us to identify this critical issue that should be carefully considered in future studies building consensus asymmetry patterns beyond humans.

Although we observe no meaningful correlations in asymmetry between species, interpretation of these results should be made cautiously. Cross-species comparisons in general are not well-defined and this represents a broader challenge facing the field of neuroimaging. To find similar regions between species, we implemented a curated, atlas-based approach, however, alignment of templates between species using white-matter estimation maps^103^ or a common transcriptional space^104^ may provide an improvement. Nonetheless, our results align with a previous study which did not observe a major cross-species signature of asymmetry between humans and chimpanzees^25^. Similarly, a study of male-female differences in humans and mice observed weak cross-species agreement, despite robust sex differences in both species^77^. Going forward, additional methods should be explored to further interrogate the question of cross-species comparisons, but at present, major consistency across species seems absent.

As brain-wide mapping studies become easier to perform in small vertebrates, we encourage authors to publish their raw images in open-source repositories like the brain image library^105^ or Zenodo.org, along with documentation to identify the left and right hemispheres. The presence in mice of a brain-wide signature of anatomical asymmetry will further enable research into the question of developmental and cellular underpinnings of asymmetry in humans, and publicly available datasets like those used in our study will be invaluable toward this effort.

Overall, we find extensive anatomical asymmetry in the adult mouse brain, manifesting as a brain-wide, anterior-posterior pattern. Our work provides a foundation for future studies to investigate the microscopic basis of this cerebral petalia-like effect in mice.

## Materials and Methods

### Overall Morphometry Analysis

Our data collection was performed using a series of python, Fiji, and R scripts. Images for each animal were first gathered into a z-stack (if not already) and converted to .tif file format from one of various starting formats (.jpg, .Jp2, .nii, .mnc, .tif). From here, each z-stack was resampled from its original resolution to 50 µm isotropic resolution. MRI samples were skull-stripped followed by N4 bias correction. We then performed image registration (rigid, affine, and non-linear) to align the sample images toward a population-average template (sample image as the moving image; reference template as the fixed image). All templates and atlases were completely left-right symmetrical. From these registrations, we collected the Jacobian determinant field and measured its average value within each CCFv3 atlas region for each sample. The Jacobian determinant field represents how much the local volume changes at each voxel when transforming one image to align with another. For example, field values of 1.10 or 0.85 at 2 different voxels in the template image indicate that the corresponding volumes in the sample image were 10% larger or 15% smaller respectively than they are in the template image. After calculating average field values for each regions, we then scaled the average field value by the physical size of that region in the original atlas to obtain a volume measurement in µm^3^. Extensive details about each cohort, registration settings, processing steps, additional analyses, left-right orientation, and more are provided in **Supplemental Materials and Methods**.

## Supporting information

Supplemental Dataset 1 -- 6-cohort average mouse brain volume asymmetry_ants

## Acknowledgments

We thank numerous members of the neuroimaging community for their generous feedback throughout this project. In this context, we would like to acknowledge Doug Greve, Elisa Guma, Stefan Klein, Yael Balbastre, and Clyde Francks. We would also like to thank individuals who provided critical information related to the left-right orientation or demographics of different cohorts: Gabriel Devenyi, John Sled, Rasmus Birn, Jonathan Oler, Junichi Hata, and Matthew Rushworth. We are thankful to Amit Zeisel for sharing supplemental data from their publication (Elkind et al. 2023^74^). We thank the Harvard Research Computing department for providing early access to the O2portal. We also thank members of the Pourquie lab for thoughtful feedback on the data and manuscript throughout the project. Finally, we are thankful for funding which this project received from a Harvard Brain Science Initiative Bipolar Disorder Seed Grant.

