## Supplemental Figures and Methods for "Left-Right Brain-Wide Asymmetry of Neuroanatomy in the Mouse Brain"

### Figure S1. Highly Significant Left-Right Volume Asymmetries in the Adult Mouse Brain.

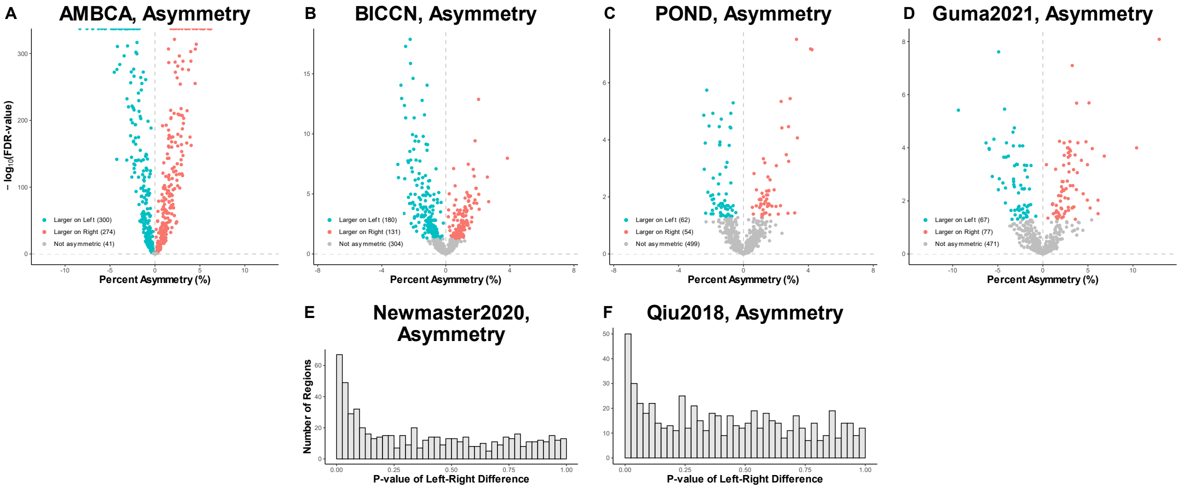


**Figure S1. A-D)** Volcano plots of significantly asymmetric brain regions for the 4 largest cohorts. **E-F)** P-value histograms of the 2 smaller cohorts for significance of each region's asymmetry. 1 sample t-test applied in all cohorts to calculate significance of a regional asymmetry from 0. FDR correction is used in A-D)**.** Nominal P-values are used for E-F).

#### Figure S2. Asymmetries Can Be Detected at 80% Power in a New Cohort with 16 Animals.

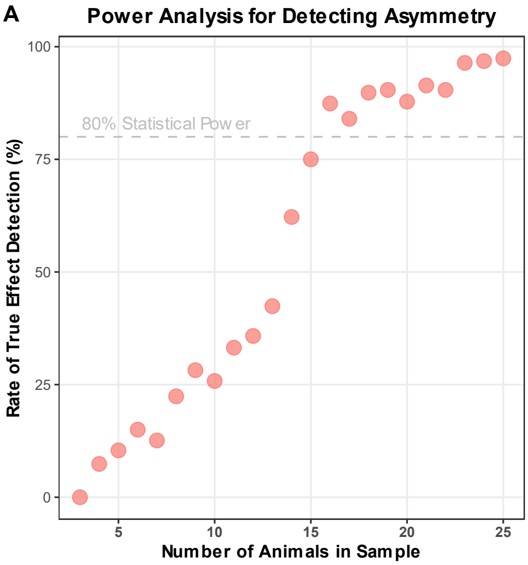


**Figure S2. A)** The percentage of samples drawn from Lopes2023 animals in which the 6-cohort asymmetry pattern can be detected successfully (Y-axis). The X-axis represents the size of the sample drawn, with a sample of that size drawn 500 times for each value on the X-axis (sampling without replacement was used). The true effect was considered successfully detected if the top asymmetries from the 6-cohort average could significantly predict the pattern of asymmetry in the correct direction in the drawn sample of animals (same as in **Fig. 1G**, but using a subset of the total Lopes2023 cohort). The 6-cohort average predicted the asymmetry pattern in 75% of samples consisting of 15 animals and in 87% of samples consisting of 16 animals.

### Figure S3. Highly Significant Asymmetries in Published Study By Independent Group.

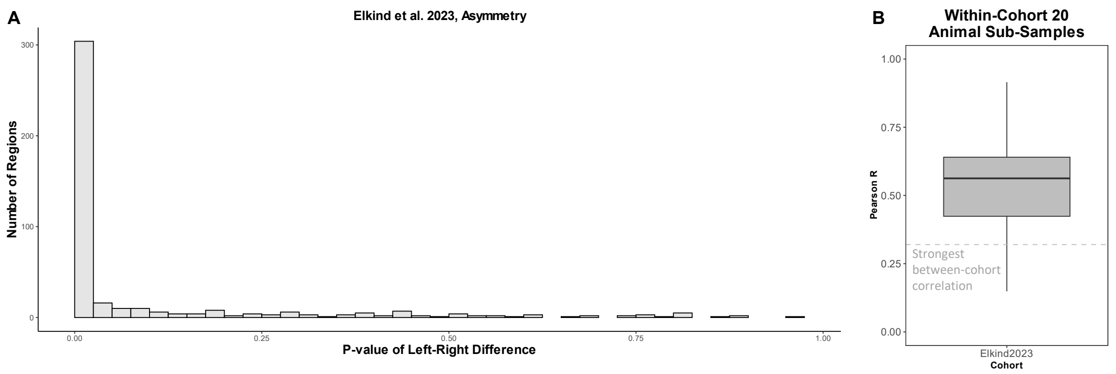


**Figure S3. A)** P-value histogram of asymmetry for the Elkind et al. 2023 data showing strong enrichment for significant asymmetries, similar to our own analysis of the mouse cohorts in this study. **B)** Distribution of Pearson R values from sub-sampling 20 animals from the Elkind et al. 2023 data and comparing the sub-sampled asymmetry pattern with their full analysis of 507 animals. Dashed line is largest non-diagonal value in **Fig. 1B**.

#### Figure S4. Analysis of Left-Right Flipped or Symmetrical MRI Images.

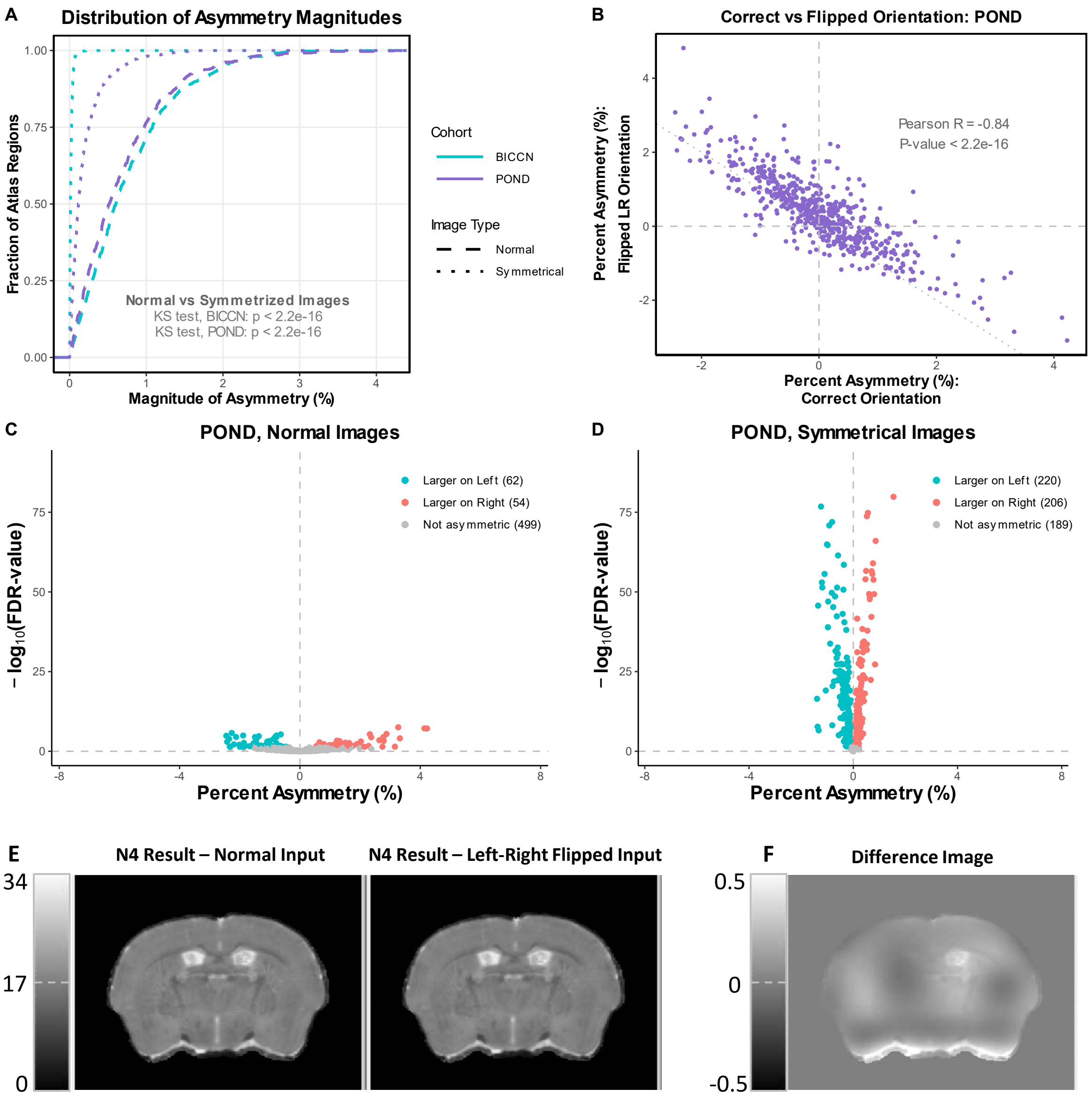


**Fig S4. A)** Cumulative density distributions for the BICCN and POND asymmetries when analyzed as the normal (non-symmetrized) images or as symmetrical images. **B)** Comparison of POND region-wise asymmetry pattern between original and left-right reversed images. Each point is a brain region where the value is the mean asymmetry for that region in each analysis. **C)** and **D)** Volcano plot of asymmetry for POND cohort where each point is a brain region. **C)** Analysis using normal images. **D)** Analysis using synthetically symmetrized samples. **E)** Example of N4 bias-corrected images generated from normal sample (left) or identical but flipped input image (right). The right image was flipped back for display purposes and for calculating the Difference Image in F). **F)** Image showing the difference between the 2 images in E). Note that the upper display limit for this "Difference Image" is 0.5, but values in this image are as large as +1.13. If outputs in E) were equivalent, then image in F) would have been all 0's.

### Figure S5. Consistency Across Registration Software For Each Cohort.

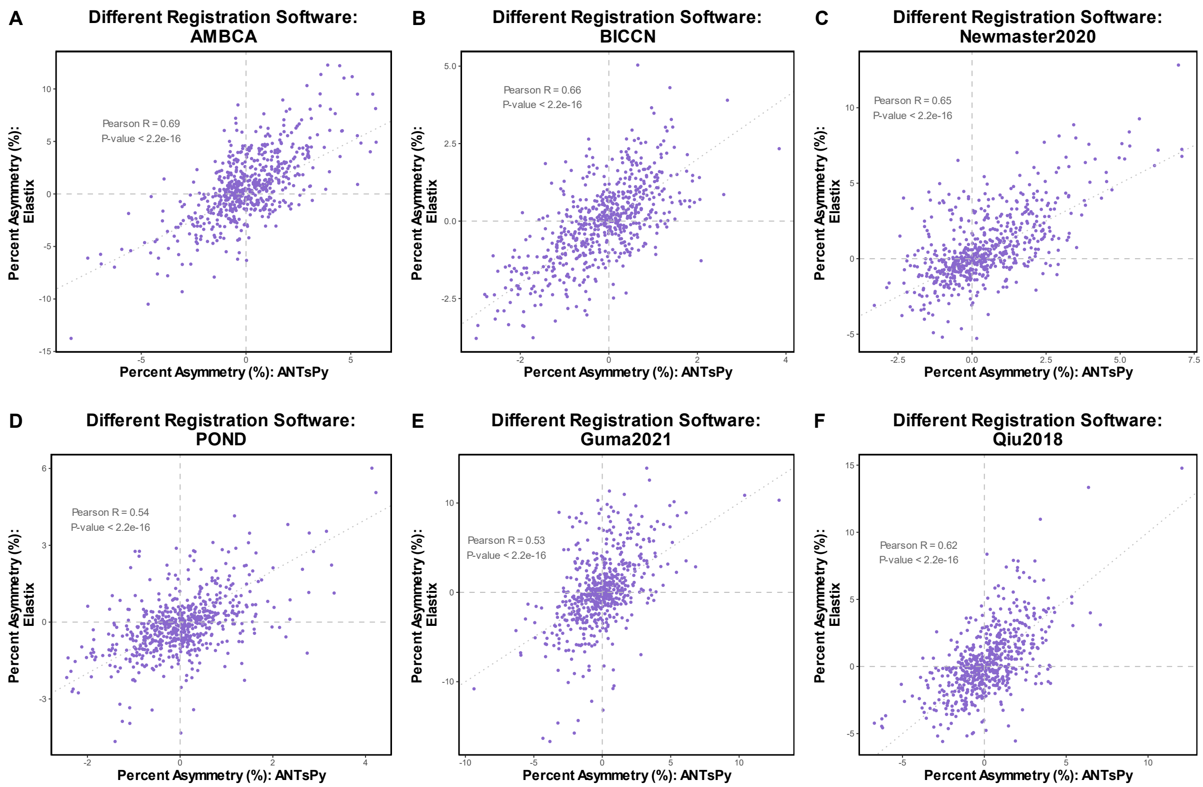


**Fig S5. A-F)** Comparison of cohort-specific region-wise asymmetry pattern generated from 2 different registration software: ANTsPy (X-axis) and Elastix (Y-axis). Each point is a brain region and the value is the average asymmetry across all animals in that cohort for each analysis.

#### Figure S6. Functional Asymmetries in the Adult Mouse Brain.

**A** **Functionally Asymmetric Brain Regions**

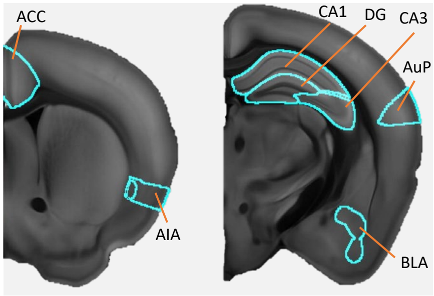


**B** **Functionally Asymmetric Brain Regions in Mice**

- **Primary Auditory Cortex** (AuP) – Response to vocalization (left), response to pitch sweeps (right)
- **Hippocampal CA3/CA1** ($CA3_{L\ vs\ R}$ -> CA1) asymmetric LTP, synapse morphology, and receptor composition; long-term spatial memory (left)
- **Dentate Gyrus** (DG) **–** Spatial representation (left), context generalization (right)
- **Anterior Insular Area** (AIA) – Response to aversive stimuli (right)
- **Basolateral Amygdala** (BLA) & **Anterior Cingulate Cortex** (ACC) – Response to observational / contextual Fear (Right for both)

**C** **Functionally Asymmetric Regions: Leave-one-out Cross-Validation**

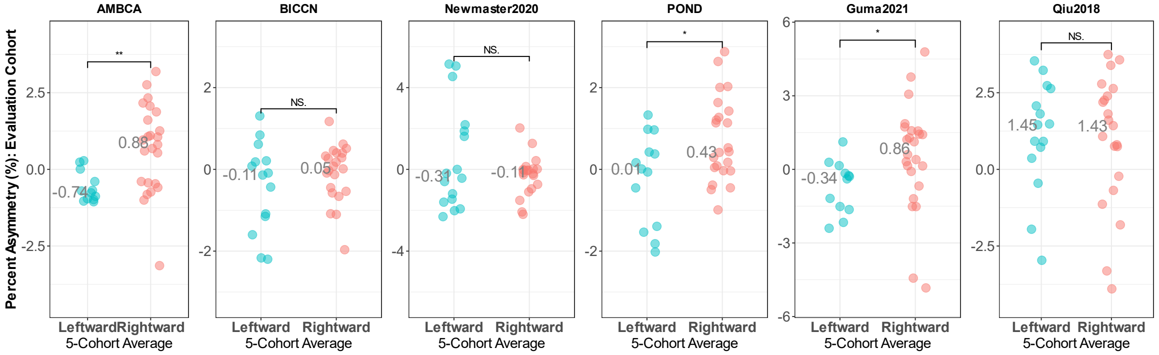


**Figure S6. A)** Areas in the mouse brain with strong evidence of functional asymmetries. **B)** List of brain regions in A) and their associated functional asymmetries. References are included in **Supplemental Table 2**. **C)** Leave-one-out Cross-validation for asymmetry using only brain regions from areas of the brain with strong evidence for functional asymmetries. Statistics are Wilcoxon rank-sum test. * is $p < 0.05$; ** is $p < 0.01$.

#### Figure S7. Additional Analysis for Anterior-Posterior Volume Asymmetry Trend.

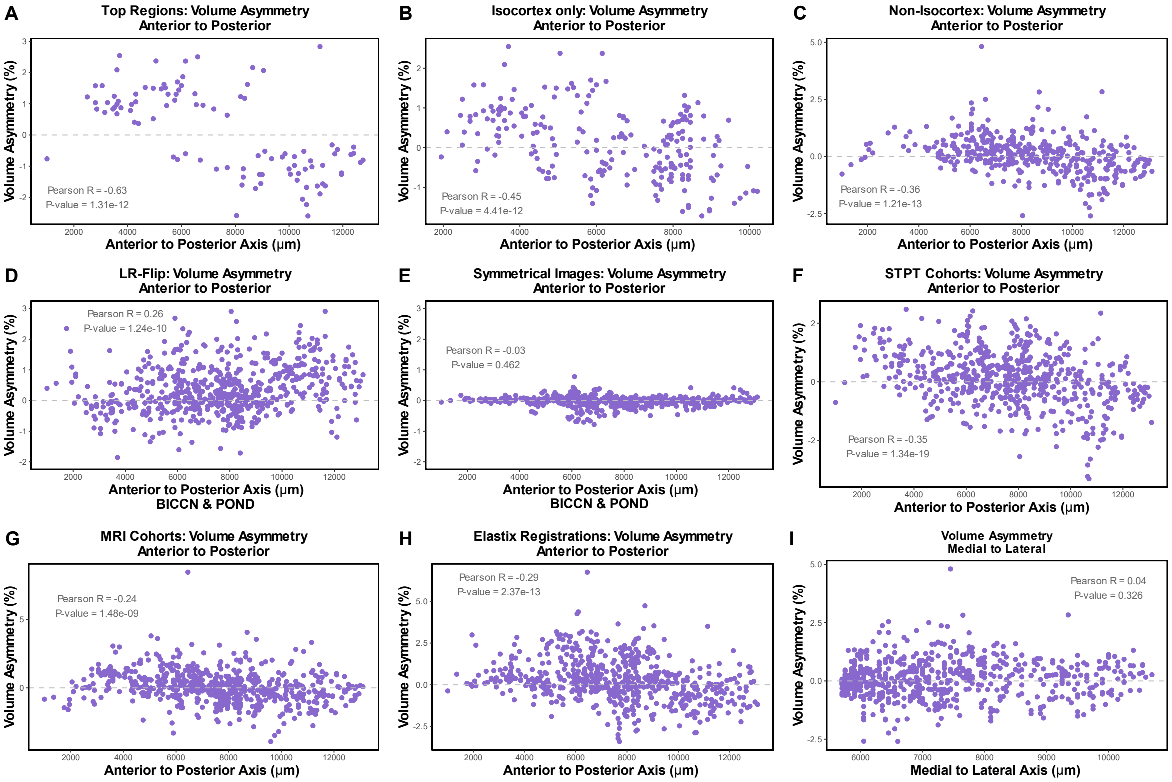


**Figure S7. A-C)** Average volume asymmetry derived from the 6-cohort average for **A)** top 100 regions, **B)** isocortex only, **C)** non-isocortex, plotted against the anterior-posterior position of that region in the atlas. **D-E)** Average volume asymmetry for all regions derived from the BICCN and POND datasets when analyzed in the **D)** Left-right flipped orientation or **E)** when symmetrized, plotted against the anterior-posterior position of that region in the atlas. **F-G)** Image modality-specific asymmetry patterns for all regions from **F)** 3 STPT cohorts (AMBCA, BICCN, Newmaster 2020) or **G)** 3 MRI cohorts (POND, Guma2021, Qiu2018), plotted against the anterior-posterior position of that region in the atlas. **H)** Average volume asymmetry derived from the 6-cohort average for all regions when performing registrations using Elastix, plotted against the anterior-posterior position of that region in the atlas. **I)** Average volume asymmetry derived from the 6-cohort average for all regions, plotted against the medial-lateral position of that region in the atlas.

**Figure S8. Anterior-Posterior Volume Asymmetry Trend is Stronger in 6-cohort average than Individual Cohorts.**

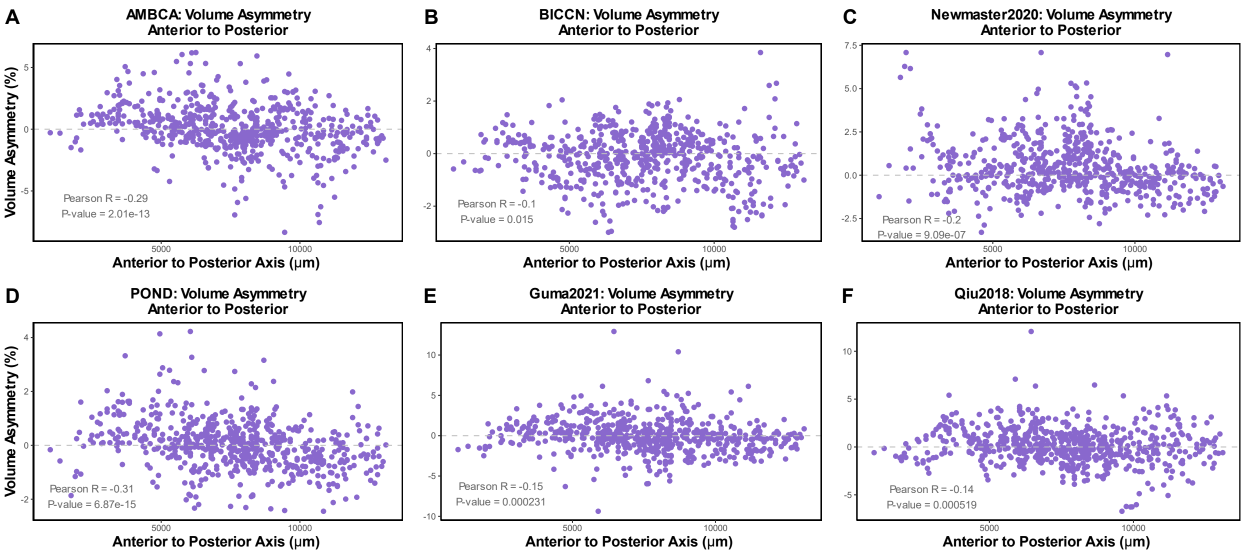


**Fig S8. A-F)** Cohort-specific volume asymmetry patterns for all regions, plotted against the anterior-posterior position of that region in the atlas. The name of the cohort from which the data is generated is specified in the graph title.

#### Figure S9. Volume Asymmetry in NHPs and Humans

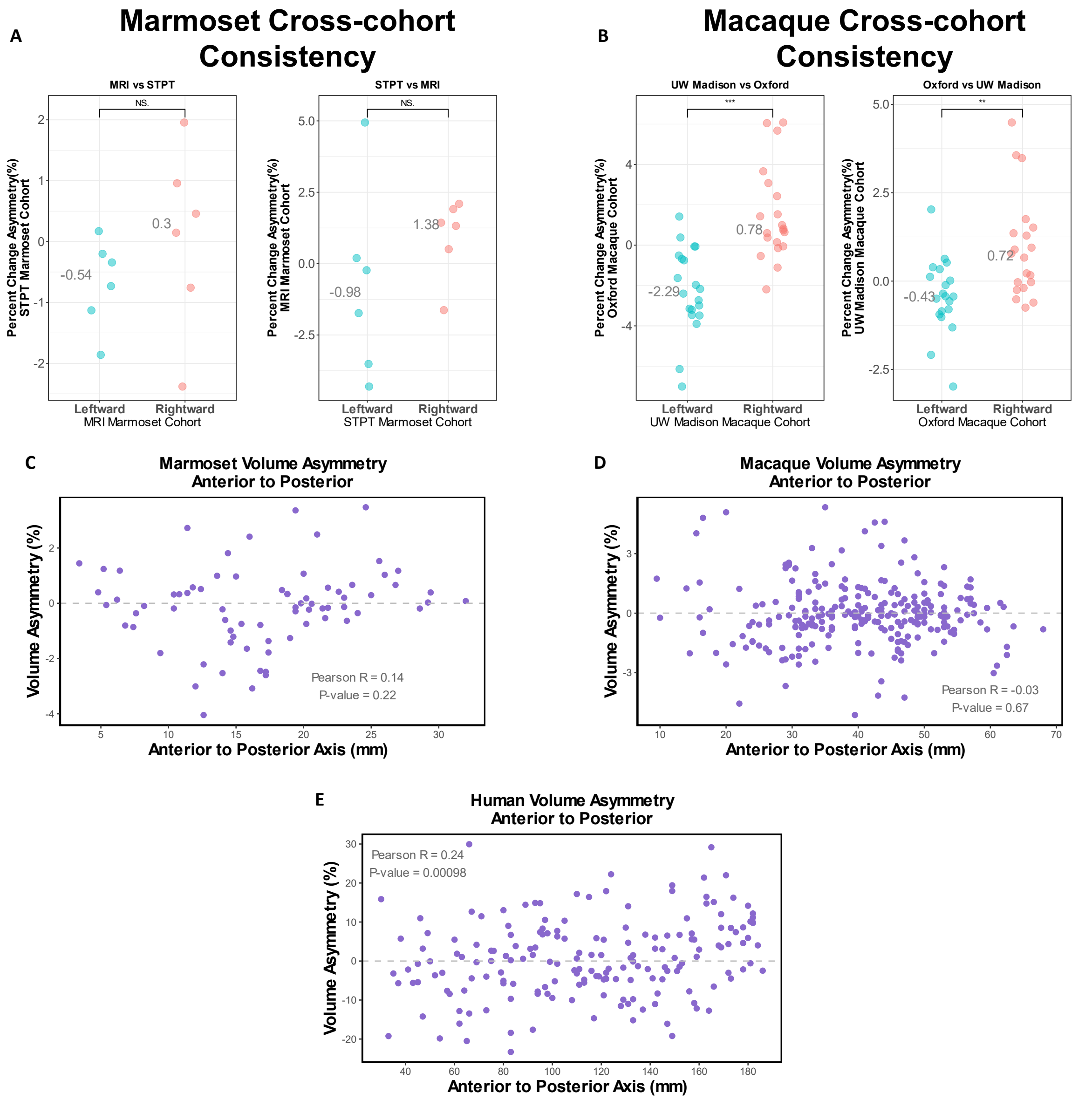


**Figure S9. A)** Cross-cohort consistency analysis between the 2 marmoset cohorts. **B)** Cross-cohort consistency between the 2 macaque cohorts. The number of regions used for A) and B) are the same proportion as the number of regions used for mice (50 leftward and 50 rightward out of 615 total; 8% of all regions for each column). **C-E)** Plotting volume asymmetry for each species across the anterior-posterior axis for **C)** Marmosets, **D)** Macaques, **E)** Humans. The average asymmetry for all cohorts of each species was used for Y-axis while the atlas 3D centroid of each region was used for the X-axis.

#### Figure S10. Cross-cohort consistency for Volume asymmetry in Humans using surface-based approach

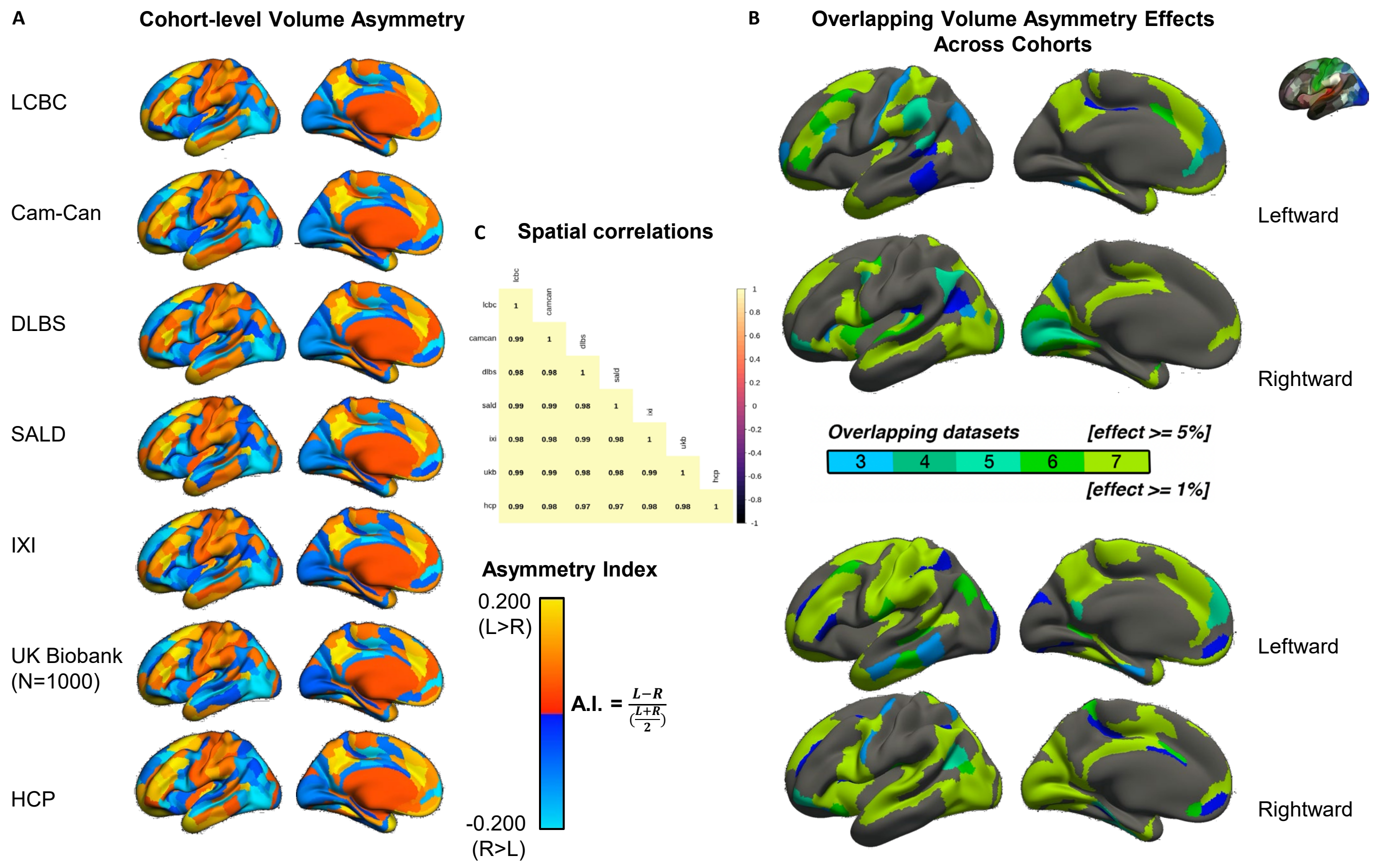


**Figure S10. A)** 3D renderings of volume asymmetry in each cohort of human subjects. Note that the metric here is laterality index instead of percent volume asymmetry. **B)** 3D renderings showing regions of interest with overlapping effects thresholded at $\geq$ 5% (top) or $\geq$ 1% (bottom). C) Pearson correlation matrix of volume asymmetry between all 7 cohorts.

#### Supplemental Table 1. Description of Mouse Cohorts.

| **Name of Cohort, Acquisition Source, and Associated Paper.** | **Image Modality** | **Number subjects** | **Strains** | **Sex Balance** | **Original Image Resolution and Initial File Format** | **Context of Dataset** | **Age** | **Notes** |
|---|---|---|---|---|---|---|---|---|
| **AMBCA**: Allen Mouse brain Connectivity Atlas.<br><br>Acquired from Allen Institute API.<br>Oh et al. 2014[106] | Serial 2-photon tomography (STPT) | 2,992 | C57Bl6/J, CD1, FVB, 129, C3H, and hybrids of these strains, as well as numerous cre recombinase strains. | 1796 male, 1196 female | Obtained .jpg files at 22.5 µm x 22.5 µm x 100 µm | Fluorescence Tracer Injections to map connectivity between brain regions | Almost exclusively P77 | Used red channel from RGB images. Acquired from Allen SDK. |
| **BICCN:** BRAIN Initiative Cell Consensus Network<br><br>Acquired from https://www.brainimagelibrary.org/<br>Hawrylycz et al. 2023[107] | STPT | 234 | 27 different fluorescent reporter strains | 118 male, 116 female | Obtained .jp2 files at 1 µm x 1 µm x 50* µm. | Mapping of cell populations in adult mouse brain and sex differences | *Not available* | *Used only every other Z plane for efficiency, so actual Z resolution is 100 µm.<br><br>Excluded "Emx1_GFP_*", "Slc17a6_GFP_*", "Snap25_GFP_*", and "Vglut1_GFP_*" subjects (39 in total) as registration for hippocampal structures was grossly inaccurate due to major intensity differences with CCFv3 template. |
| **Newmaster2020:** Newmaster et al. 2020[108]<br><br>Acquired from https://www.brainimagelibrary.org/ | STPT | 23 adults | Oxytocin transgenic reporter mice | 12 male, 11 female | Obtained .tif files at 1 x µm x 1 µm x 50* µm. | Oxytocin receptor mapping over development. | P56 | Some tissue defects, optical defects. *Used only every other Z plane for efficiency, so actual Z resolution is 100 µm. |
| **POND:** Province of Ontario Neurodevelopmental network<br><br>Acquired from https://www.braincode.ca/content/open-data-releases after requesting access.<br>Ellegood et al., 2015[109]<br>Most of these mice were bred and perfused in different labs, then shipped to the Lerch lab for imaging. | T2w *Ex Vivo* MRI | 207 wild-type | Wild-type littermates from many different autism genetic strains. Strains on C57BL6/J, C57BL6/N, FVB, 129, or other backgrounds. | 140 male, 67 female | All Isotropic: either at 32 µm, 40 µm, or 56 µm resolution. Obtained .mnc files. | MRI Imaging of autism spectrum disorder mice | P58-P98 | Original data provided by POND network had issue in which 173 wild-type scans were duplicates of other scans. This has since been corrected by the POND network but I never went back and included those mice. I use only the original 207 wild-type mice which were provided correctly upon initial download. |
| **Guma2021:** Guma et al. 2021[110]<br><br>Acquired from https://zenodo.org/records/5156508 | T1w MEMRI *In Vivo* MRI | 41 saline-injected mice | C57BL/6J | 21 male, 20 female | Obtained .mnc files at 100 µm isotropic. | Impact of gestational immune activation | Longitudinal imaging: P21, P60, and P90, | P60 and P90 scans averaged for "adult" time-points |
| **Qiu2018:** Qiu et al. 2018[76]<br><br>Acquired from Jacob Ellegood and Jason Lerch.<br><br>These mice were bred and raised by the Lerch lab. | T1w MEMRI *In Vivo* MRI | 23 | C57BL/6J | P65: 9 male, 10 female<br><br>P36: 2 male, 2 female | Obtained .mnc files at 90 µm isotropic. | Plausibility of T1w MEMRI for studying postnatal development in mice | 19 P65 mice and 4 P36 mice. | Low signal-noise ratio is apparent in many samples. Prior to any analysis, five P65 mice were dropped due to low image quality and replaced with four P36 scans of different animals. |
| **Lopes2023:** Lopes et al. 2023 (Elife)<br>Acquired from https://osf.io/m7gpd/<br>***Reserved for generalizability*** | T2w *Ex Vivo* MRI | 48 | C57BL/6JRj | All female | Obtained .nii.gz files at 100 µm isotropic. | Impact of environmental enrichment on brain volume | 17 weeks old | |

**Supplemental Table 2. Papers Showing Functional Asymmetry in the Mouse Brain.**

| Brain Region | Publication Title | Authors, year, journal | Paper Link on PubMed |
|---|---|---|---|
| **Left Primary Auditory Cortex** | Left hemisphere advantage in the mouse brain for recognizing ultrasonic communication calls | Gunter Ehret, 1987, Nature | https://pubmed.ncbi.nlm.nih.gov/3808021/ |
| **Left Primary Auditory Cortex** | Oxytocin enables maternal behavior by balancing cortical inhibition | Marlin et al. 2015, Nature | https://pubmed.ncbi.nlm.nih.gov/25874674/ |
| **Left Primary Auditory Cortex** | Bilateral widefield calcium imaging reveals circuit asymmetries and lateralized functional activation of the mouse auditory cortex | Calhoun et al. 2023, PNAS | https://pubmed.ncbi.nlm.nih.gov/37459544/ |
| **Left and Right Primary Auditory Cortex** | Circuit asymmetries underlie functional lateralization in the mouse auditory cortex | Levy et al. 2019, Nature Communications | https://pubmed.ncbi.nlm.nih.gov/31239458/ |
| **Left and Right Primary Auditory Cortex** | Differences in temporal processing speeds between the right and left auditory cortex reflect the strength of recurrent synaptic connectivity | Neophytou et al., 2022, PLoS Biology | https://pubmed.ncbi.nlm.nih.gov/36269764/ |
| **Hippocampus (CA3-CA1)** | Asymmetrical allocation of NMDA receptor epsilon2 subunits in hippocampal circuitry | Kawakami et al., 2003, Science | https://pubmed.ncbi.nlm.nih.gov/12738868/ |
| **Hippocampus (CA3-CA1)** | Target-cell-specific left-right asymmetry of NMDA receptor content in schaffer collateral synapses in epsilon1/NR2A knock-out mice | Wu et al., 2005, J. Neurosci | https://pubmed.ncbi.nlm.nih.gov/16207881/ |
| **Hippocampus (CA3-CA1)** | Left-right asymmetry of the hippocampal synapses with differential subunit allocation of glutamate receptors | Shinohara et al., 2008, PNAS | https://pubmed.ncbi.nlm.nih.gov/19052236/ |
| **Hippocampus (CA3-CA1)** | Left-right asymmetry defect in the hippocampal circuitry impairs spatial learning and working memory in iv mice | Goto et al., 2010, PLoS One | https://pubmed.ncbi.nlm.nih.gov/21103351/ |
| **Hippocampus (CA3-CA1)** | Hemisphere-specific optogenetic stimulation reveals left-right asymmetry of hippocampal plasticity | Kohl et al., 2011, Nature Neuroscience | https://pubmed.ncbi.nlm.nih.gov/21946328/ |
| **Hippocampus (CA3-CA1)** | Left-right dissociation of hippocampal memory processes in mice | Shipton et al., 2014, PNAS | https://pubmed.ncbi.nlm.nih.gov/25246561/ |
| **Hippocampus (CA3-CA1)** | PirB regulates asymmetries in hippocampal circuitry | Ukai et al., 2017, PLoS One | https://pubmed.ncbi.nlm.nih.gov/28594961/ |
| **Hippocampus (CA3-CA1)** | The lateralization of left hippocampal CA3 during the retrieval of spatial working memory | Song et al., 2020, Nature Communications | https://pubmed.ncbi.nlm.nih.gov/32518226/ |
| **Hippocampus (CA3-CA1)** | An emergent neural coactivity code for dynamic memory | El-Gaby et al., 2021, Nature Neuroscience | https://pubmed.ncbi.nlm.nih.gov/33782620/ |
| **Hippocampus (Dentate Gyrus)** | Hemisphere-specific spatial representation by hippocampal granule cells | Cholvin and Bartos, 2022, Nature Communications | https://pubmed.ncbi.nlm.nih.gov/36266288/ |
| **Right Insular Cortex** | The anterior insular cortex unilaterally controls feeding in response to aversive visceral stimuli in mice | Wu et al., 2020, Nature Communications | https://pubmed.ncbi.nlm.nih.gov/32005806/ |
| **Right Insular Cortex and Basolateral Amygdala** | Glutamatergic synapses from the insular cortex to the basolateral amygdala encode observational pain | Zhang et al., 2022, Neuron | https://pubmed.ncbi.nlm.nih.gov/35443154/ |
| **Right Basolateral amygdala** | Contextual fear conditioning is associated with lateralized expression of the immediate early gene c-fos in the central and basolateral amygdalar nuclei | Scicli et al., 2004, Behavioral Neuroscience | https://pubmed.ncbi.nlm.nih.gov/14979778/ |
| **Right Basolateral amygdala and right anterior cingulate cortex** | Hemispherically lateralized rhythmic oscillations in the cingulate-amygdala circuit drive affective empathy in mice | Kim et al., 2023, Neuron | https://pubmed.ncbi.nlm.nih.gov/36460007/ |
| **Right Anterior Cingulate Cortex** | Lateralization of observational fear learning at the cortical but not thalamic level in mice | Kim et al., 2012, PNAS | https://pubmed.ncbi.nlm.nih.gov/22949656/ |

#### Supplemental Table 3. Description of Human and Non-Human Primate Cohorts.

| Animal/Species | Name | Image Modality | Number subjects | Sex Balance | Original Image Resolution | Context of Dataset | Age | Notes |
|---|---|---|---|---|---|---|---|---|
| **Humans**<br>(Homo sapiens)<br><br>Acquired from<br>https://openneuro.org/datasets/ds004215/versions/1.0.0 | **NIMH**: National Institute of Mental Health.<br>Nugent et al. 2022[111] | T1w *In Vivo* MRI | 155 | 53 male,<br>102 female | 1 mm isotropic | Assemble dataset of healthy adult volunteers. | 18 – 72 years old | |
| **Humans**<br>(Homo sapiens)<br><br>Acquired from<br>https://openneuro.org/datasets/ds004173/versions/1.0.2 | **MRART:** Movement-related artefacts.<br>Nárai et al. 2022[112] | T1w *In Vivo* MRI | 148 movement-free scans | 53 male,<br>95 female | 1 mm isotropic | Study effects of subject movement during MRI scan on image quality. | 18 – 75 years old | |
| **Humans**<br>(Homo sapiens)<br><br>Acquired from<br>https://brain-development.org/ixi-dataset/ | **IXI:** Information eXtraction from Images<br><br>https://brain-development.org/ixi-dataset/ | T1w *In Vivo* MRI | 619 | 277 male,<br>342 female | In-plane: 0.9375 mm x 0.9375 mm<br>Between planes: 1.2 mm | Normal, healthy subjects | 19-86 years old | |
| **Humans**<br>(Homo sapiens)<br><br>Acquired from<br>https://fcon_1000.projects.nitrc.org/indi/retro/sald.html | **SALD:** Southwest university adult lifespan dataset<br>Wei et al. 2018[113] | T1w *In Vivo* MRI | 494 | 187 male,<br>307 female | 1 mm isotropic | Largely university students and staff without history or current neurological/psychiatric disorders. No history of head trauma. | 19-80 years old | |
| **Chimpanzees**<br>(Pan troglodytes)<br><br>Acquired from Bill Hopkins at National Chimpanzee Brain Resource upon request. | **Yerkes:** Yerkes National Primate Laboratory, Atlanta, Georgia | T1w *In Vivo* MRI | 73 | 26 male,<br>47 female | Variable, between 350 µm non-isotropic and 1 mm isotropic. Many were not isotropic, with 2-fold lower Z-step than in-plane resolution, several had 4-fold lower Z-step compared to in-plane resolution. | Study of healthy, captive chimpanzees | 9 – 53 years old | |
| **Chimpanzees**<br>(Pan troglodytes)<br><br>Acquired from Bill Hopkins at National Chimpanzee Brain Resource upon request. | **Bastrop:** MD Anderson, Bastrop, Texas | T1w *In Vivo* MRI | 139 | 58 male,<br>81 female | Same as above | Study of healthy, captive chimpanzees | 8 – 51 years old | |
| **Macaques**<br>(Macaca Mulatta) | **HL** and **WNPRC:**<br>Harlow Lab and Wisconsin National Primate Center | T1w *In Vivo* MRI | 592 | 327 male,<br>265 female | 273 µm x 500 µm x 273 µm | Study of brain development in juvenile macaques | ~10 months | HL and WNPRC are |

| | | | | | | | | |
|---|---|---|---|---|---|---|---|---|
| Acquired from PRIME-de | https://fcon_1000.projects.nitrc.org/indi/PRIME/uwmadison.html | | | | | | old to 4.5 years old. | Adjacent facilities. |
| **Macaques** (Macaca mulatta) Acquired from PRIME-de | **Oxford**: Wellcome center for integrative neuroimaging Noon et al. 2014[114] | T1w *In Vivo* MRI | 20 | All male | 500 µm isotropic | unknown | 2.4-6.7 years old | |
| **Marmosets** (Callithrix jacchus) Acquired from Brain/MINDS | **NA216:** Brain/MINDS Marmoset Brain MRI Database Hata et al. 2023[115] | T2w *In Vivo* MRI | 216 | 93 male, 132 female | 270 µm XY resolution 540 µm Z resolution | Generation of public marmoset MRI dataset | 1 – 12.5 years | |
| **Marmosets** (Callithrix jacchus) Acquired from Brain/MINDS | **BMCA:** Brain/MINDS Marmoset Connectivity Atlas Skibbe et al. 2023[116] | STPT | 44 | 18 male, 26 female | 1.3 µm XY resolution 50 µm Z resolution | Marmoset prefrontal cortex connectivity mapping | 2.3 – 10.8 years | |

#### Supplemental Materials and Methods

**Python and Computing Environment.** All code for performing our analysis is available at https://osf.io/wkhxs/. Our analysis relied heavily on ANTsPy[71] and ClearMap2[117] functions. We provide a version-stable environment.yml file to replicate our environment. We performed our analysis on the O2 high performance computing cluster at Harvard Medical School, specifically using the O2portal to enable interactive analysis and visualization. O2 is a shared computational resource across Harvard Medical School containing over 390 compute nodes, 12,000 compute cores, and more than 100TB of memory.

**Conversion of Various File Formats to .tif and Resampling.** Due to the challenges of working with a large number of samples (AMBCA), extremely large single-plane images (BICCN, Newmaster et al. 2020), and varying file formats, we used slightly different steps to convert the images to .tif and resample each cohort to 50 µm isotropic resolution. STPT cohorts were obtained as series of 2D .Jpg, .Jp2, and .tif images, where each image consisted of 1 coronal plane. MRI cohorts were obtained either as 3D .mnc or 3D .nii files (**Table S1**).

For the AMBCA, we downloaded images from the Allen Institute API which were approximately 22.4 µm XY resolution (6X down-sampled), with 100 µm between planes. Each AMBCA animal's RGB images were loaded as an image sequence in Fiji and resampled to 50 µm isotropic resolution, keeping only the red channel. All other cohorts were grey-scale images. For the BICCN and Newmaster et al. 2020 cohorts, we obtained the original images at ~1 µm XY resolution with 50 µm between planes. We then down-sampled the dorsal-ventral axis of the images using opencv2 in python before loading the images in PyImageJ (python implementation of Fiji) for the rest of the resampling. For computational efficiency, we used every 2$^{nd}$ image plane in these cohorts, initially resulting in 100 µm Z spacing (same as full AMBCA) which we then up-sampled to 50 µm spacing for analysis. For the MRI cohorts, we converted the initial .mnc files first to .nii using Mnc-Toolkit V2, and then used nibabel in python to read the images as numpy arrays and save them as .tif files. MRI scans were resampled using PyImageJ to 50 µm isotropic resolution from their various starting resolutions. All information pertaining to left-right orientation of the images was considered at the end of the pipeline (see Left-right orientation section).

**Skull-stripping and N4 Bias Correction.** For MRI cohorts, we performed skull-stripping using ANTsPy. We did this by performing 1 or 2 initial rigid registrations to first align the samples into a common pose. Then we linearly and non-linearly registered a template image with non-brain tissue present onto each rigidly-aligned sample brain. Using this resulting transformation, we transformed the atlas labels into the sample space. The area of the sample covered by the warped atlas was used to define the brain in each sample. We then binarized this area and expanded it slightly to provide a small buffer around the brain without reaching the skull or doing so minimally. All pixels in the sample image beyond this space were set to 0, thus removing the surrounding tissue. A similar buffer was left when skull-stripping templates that we generated in-house. Registration settings for the nonlinear portions of skull-stripping were held constant across cohorts for a given species but varied slightly across species. These skull-stripped images were then bias-corrected using the ANTsPy N4 algorithm with the default settings.

**Jacobian-generating Registrations.** Our final registrations from which we obtained the Jacobian determinant fields were performed using ANTsPy[71] or Elastix[72] (when comparing the 2 software). The registration settings of each software were held constant for all mouse cohorts and were modified only slightly in ANTsPy for the other species (constant settings within each species). For ANTsPy registrations, we first rigidly aligned each sample as the moving image into the template space (fixed image; used mutual information metric). From there, we used an affine (mutual information metric) registration followed by symmetric normalization (normalized cross correlation metric). The Jacobian determinant fields were extracted from these registrations for analysis. The calls we made to the python ANTsPy registration command are below:

**Rigid alignment registration call:**

*All Species:* **ants.registration**(fixed=fi, moving=mi, type_of_transform = 'Rigid', verbose = True)

**Affine and Nonlinear registration call:**

*Mice:* **ants.registration**(fixed=fi, moving=mi, type_of_transform = 'SyN',syn_metric='CC', syn_sampling = 3, verbose = True, flow_sigma = 2, total_sigma= 0.3, grad_step =0.1, reg_iterations=(100,100,50,25), write_composite_transform=False)

*Marmosets*, *Macaques, Chimpanzees, and Humans:* **ants.registration**(fixed=fi, moving=mi, type_of_transform = 'SyN',syn_metric='CC', syn_sampling = 3, verbose = True, flow_sigma = 1, total_sigma = 0.1, grad_step = 0.3, reg_iterations=(100,100,50,25), write_composite_transform=False)

- "fi" and "mi" are each a 3D image series loaded into memory as an ANTsPy image.
- "fi" is the template (CCFv3) while "mi" is the sample.
- An affine registration is performed when specifying the "SyN" transform parameter and has a default metric of "mattes" (mattes mutual information).

For registrations performed in Elastix, we similarly started with a rigid registration ("AdvancedNormalizedCorrelation" metric), followed by an affine ("AdvancedNormalized Correlation" metric) and bspline registration ("AdvancedMattesMutualInformation" metric with "TransformBendingEnergyPenalty"). Too many parameters exist in Elastix to describe the settings here, however the full parameter files are available along with the rest of the code for performing all the analysis at https://osf.io/wkhxs/. To summarize the Elastix settings, the rigid and affine parameter files were modified versions of Par0025rigid.txt and Par0025affine.txt files from the Elastix parameter model zoo. The bspline registration was a modified version of the align_bspline.txt parameter file obtained with the ClearMap2 software package. Briefly, our nonlinear registration in Elastix used 4 resolutions of 5000 iterations each, with a grid spacing schedule of 8,4,2,1. The final grid spacing resolution in voxels was 4. The weight of Metric0 (AdvancedMattesMutualInformation) was 1 while Metric1 (TransformBendingEnergy Penalty) was 50 to prevent negative jacobians. The default "SmoothingImagePyramid" was used for the fixed and moving images.

**Templates used (Volumetric registrations).** For mouse STPT cohorts, we utilized the CCFv3 population average template and atlas for registration. For the Guma2021 and Qiu2018 cohorts, we used modality-specific average templates which we generated in-house using these cohorts (using SimpleElastix) and which we subsequently aligned to the CCFv3 space. For the POND cohort, we used the previously generated template provided with the images which we aligned to CCFv3 space and symmetrized. For cross-species analysis, the Garin et al. 2022 paper used the DSURQE template and atlas for the mouse portion. As such, we warped the DSURQE template and atlas images into CCFv3 space and re-calculated the mean jacobians for each parcel. For humans, we used the "MNI ICBM152 nlin asym 09c" template (https://www.bic.mni.mcgill.ca/ServicesAtlases/ICBM152NLin2009) which we symmetrized, along with the Glasser HCP-extension atlas[118]**.** For chimpanzees, we used the DaVi130 atlas and associated template. Macaque registrations used the NMTv2.0 template with associated CHARM, SARM, and D99 atlases. Our final macaque parcellations used the level 5 CHARM and SARM annotations. The STPT Marmosets used the Brain MINDS Connectivity Resource V2 STPT template while the MRI marmosets used the “NA216 T2WI in STPT space” template. We symmetrized all templates and atlases used (see below).

**Symmetrizing of Templates.** All templates, atlases, masks, or other derivative images used in this project were made absolutely symmetrical, regardless of whether the image was previously generated by another group or produced in-house. We did this by selecting one hemisphere (usually the right) and duplicating it across the midline. When generating in-house templates, we often did this multiple times throughout the process and again with the final utilized files. We did this simultaneously with any masks or atlases associated with the template to ensure symmetry in all of the relevant files. We checked that this process produced symmetrical images by reflecting the image over the mid-point of the X axis and subtracting the flipped image from the original image. Symmetrical images produce all 0’s in this context, which we always observed.

**Quality Control During Image Processing.** Quality control was performed after N4 normalization, skull-stripping, and our final Jacobian-generating registrations by inspecting a montage of 5 resulting image planes from each animal spanning the full brain, either overlayed against the registration template (skull-stripping and final registration) or by themselves (N4). This was done using Fiji scripts. Any animal with gross mis-registrations or other defect was noted and excluded from the analysis. Registration quality was generally high, even in the lowest signal-to-noise ratio dataset (an *in vivo* MRI cohort). One localized but recurring mis-registration was seen in the macaque cohorts, however the overall pattern of asymmetry was consistent between animals with and without this issue, thus we used both groups of animals. No other systematic issues were observed, especially in mice. For the AMBCA cohort, we inspected only the first 250 animals while we inspected every animal in other mouse cohorts.

**Tabular Analysis of Morphometry Results.** Each animal’s Jacobian determinant field was loaded into python and the average Jacobian value was measured within each annotated region. This was then saved as a .csv file and imported into Rstudio. We excluded a region from a single animal if its volume was greater than 3 standard deviation (SD) from the cohort-mean of any given structure (hemisphere-specific means). If one side of a particular region was dropped in an animal, the associated region in the opposite hemisphere was also dropped. Any animal with > 10% of all regions beyond 3 SD was excluded entirely. All samples noted to have poor registrations or negative Jacobian determinants were excluded from analysis too (as in the Quality Control section). Instances of negative Jacobians were less common in ANTsPy than Elastix (still uncommon overall), given that the ANTsPy algorithm constrains the registration generally to produce positive Jacobian determinants.

Asymmetry of each region was calculated for each animal as:

$$asymmetry = 100\% \times \frac{Right - Left}{Left}.$$

The asymmetry of each region was then averaged over all animals in the cohort to get a cohort-mean asymmetry. One-sample t-tests of the population mean were performed against 0 (symmetry) for each region, followed by FDR correction to determine number of significant regions after multiple-testing correction. We used un-adjusted p-values to rank the 50 most significant asymmetries where the left was larger and 50 most significant where the right was larger (set of “top regions”; 100 total out of 615 regions in the atlas).

These collections of leftward and rightward top asymmetries defined in one cohort were then measured in another cohort and the resulting asymmetry was tested between the 2 groups using Wilcoxon non-parametric t-tests. We report the Hodges-lehmann estimator for the difference in central tendency in our matrix figures given that the distributions of regions with leftward and rightward asymmetries had varying shapes and so the difference in means or medians is less informative.

Identical procedures were followed for evaluating sex differences but we normalized the regional volumes for each animal by the total volume of each individual animal's brain. We did this due to previous reports of a large male mouse brain volume than females, although we observed no consistent male-female global difference in brain volume. Here, we used the formula:

$$Sex\ difference = 100\% \times \frac{Males - Females}{Females}$$

to calculate significance of sex differences within a single cohort, but instead of the 1-sample paired t-test used for asymmetry, we used a 2-sample un-paired t-test between males and females. Between-cohort comparisons of sex differences still used the Wilcoxon non-parametric t-tests for generating statistics.

**Cross-validation procedure.** Leave-one-out cross-validation was performed by averaging the summary patterns of asymmetry from 5 cohorts together. We did not weight the summary patterns by the number of samples in each cohort because the strong cohort-specific effects for asymmetry makes it unlikely that the pattern of a really large cohort is more accurate than the pattern of a smaller one. In each 5-cohort average, we calculated a new average asymmetry for all 615 regions based on the mean of the 5 cohort summaries for that region. We then used a one-sample t-test on the cohort means to calculate p-values: i.e. the probability that the distribution of these 5 cohort means was different from 0. We used that p-value to again define the top asymmetries (50 leftward, 50 rightward) from this 5-cohort average and evaluate the asymmetry of these regions in the remaining $6^{th}$ cohort using a Wilcoxon non-parametric t-test. Identical procedures were again followed for calculating image modality-specific patterns or when evaluating sex differences.

**Calculation of Atlas Centroids.** Atlas centroids were obtained by first loading the 3D atlas image into python as a NumPy array. For each annotated atlas region, we then identified the coordinates of all the voxels corresponding to that region. We calculated the 3D centroid as the average coordinate of each region in X, Y, and Z from that region's identified voxels.

**Calculation of Surface Area and Cortical Thickness.** Surface area and cortical thickness were measured through a multi-step process. This can be summarized as 1) build a pial surface model and calculate its area per cortical territory (surface area), 2) build a model of the white matter-cortex interface, 3) generate a smooth Laplacian field between the pial surface model and the white matter/cortex interface model, and 4) calculate the steepest descent path through the Laplacian field from each pial surface voxel to the white matter/cortex surface. The mean length of this descent path, per cortical territory, was our cortical thickness measurement. Our algorithm is based on that of Jones et al. 2000[78] and Lerch et al. 2008[79]. Of note, we define cortical "territory" as all layers for a given cortical area collapsed into 1 label: e.g. a "territory" is "Primary motor area" which is comprised of Primary motor area, layers 1-6b.

To elaborate on our implementation, we first reversed the Jacobian-generating registration from ANTsPy to map atlas labels into sample space. We then used these sample-space annotations to reconstruct the cortex of each sample as a NumPy array, each hemisphere separately. We used SciPy edge detection to define the complete boundary of the cortex in each hemisphere, which we thresholded and then skeletonized using skeletonize_3d from skimage.morphology. This produced a thin surface completely enclosing the mouse cortex. We note that this function actually produces different skeletons depending on whether a given line curves to the left or to the right. As such, we built both hemispheres separately – temporarily reflecting the right hemisphere to the left to implement this function and then correcting this reflection after this step. We brought the two hemisphere skeletons together to create 1 cortical surface model with both hemispheres.

We next annotated this cortical surface model to indicate whether each surface voxel corresponded to the pial surface, the white-matter/cortex interface, or an interface with another structure such as the piriform cortex or the interhemispheric fissure. Generally, the pial surface was defined as any cortical surface voxel bordering "universe" voxels (voxels outside of the brain; empty space). The white-matter interface was defined as cortical surface voxels bordering white matter tracts or other select non-cortical brain tissue. A 3rd class of annotations was used to model special cortical geometry – for example, the piriform cortex, which directly borders the isocortex in a manner perpendicular to the cortical layers rather than directly beneath them as for the white matter. Numerous exceptions to this classification scheme exist which are too detailed to list directly but which were added to improve the boundary annotations. For example, we set a condition for cortical voxels bordering the olfactory bulb to be treated as pial boundary, even though they did not border the "universe". The full implementation is available in the script "8_corticalThickness_ccfv3.py" at https://osf.io/wkhxs/.

From this annotation process, the cortex surface model was output with the pial surface defined as a pixel value of 3, the white-matter interface defined as a value of 1, and the empty space within the surface set as a value of 2. The 3rd class of boundary annotations for special cortical geometry was set to a value of 4. Surface area measurements were obtained by counting all cortical surface voxels with a value of 3 per cortical territory from the CCFv3 atlas labels.

This annotated surface model was then used to generate a smooth 3D Laplacian field within it by Jacobi iteration: take the average of all 6 non-diagonal neighbors surrounding each pixel and set that as the voxel's new value:

$$new\ value_{x,y,z} = \frac{1}{6}(old\ value_{x-1,y,z} + old\ value_{x+1,y,z} + old\ value_{x,y-1,z} +$$

$$old\ value_{x,y+1,z} + old\ value_{x,y,z-1} + old\ value_{x,y,z+1})$$

The third class of annotations (value = 4) was treated as a "resistive boundary": If a non-boundary voxel (one with an initial value = 2) was neighbors with a resistive boundary voxel (initial value = 4), that resistive boundary voxel was not included in the above calculation and the denominator of $\frac{1}{6}$ was decreased accordingly ($\frac{1}{5}$ if 1 resistive voxel, $\frac{1}{4}$ if 2, etc…). Overall, this process of calculating and updating the 3D field was repeated for a maximum of 10,000 iterations until the cumulative difference between the field in 1 iteration and the next was less than $1 \times 10^{-5}$ (convergence). As the pial surface, white-matter/cortex interface, and resistive boundaries were all fixed in their values throughout this process (3,1,4 respectively), all field values fall within the interval (1,3) exclusive. An example of a converged field is shown in the left hemisphere of **Fig. 5C**.

To measure the thickness of the cortex, we then performed gradient descent through the this converged field, for all annotated pial-surface voxels, until arriving at the white-matter interface. The total length of the descent path was used as a single pial surface voxel's measure of cortical thickness, and thickness values were averaged for all voxels within each cortical territory.

**Calculation of the Cerebral Petalia.** We measured the cerebral petalia (anterior and posterior protrusion of one hemisphere with respect to the other) using an approach based on 2 recent, automated studies of the petalia[13,14]. The key concept from these studies is that when using a non-deforming registration (rigid or affine), the petalia asymmetry itself in humans may prevent an accurate registration of a subject's brain midline with the true midline of an average template. This would ultimately produce an inappropriate reference frame for measuring an anterior-posterior protrusion. Instead, previous authors defined the mid-sagittal plane (MSP) for each sample and aligned that plane to the atlas's mid-sagittal plane. We implemented a similar approach first in humans, and then for all other species.

To define the MSP in each subject, we made a custom atlas image for each species focused on the midline regions, starting from the full, regionally annotated atlas. We deleted voxels more than 5 voxels away from either side of the midline. We then cropped away the anterior and posterior ends of the atlas such that the remaining tissue was located approximately in the middle 2/3rds of the brain's AP axis. The end result is that the final, modified atlas contained tissue only along the core of the AP axis and away from the cortical poles, in the most medial (medial-lateral axis) portions of the brain. We call this image the MSP-atlas (these are included in our code/data repository).

Having previously used a full linear and non-linear registration sequence to accurately propagate the full atlas annotations into the space of each sample, we re-used that registration to map our new custom MSP-atlas onto each sample. We then performed a rigid registration (rotation, translation) to align that sample-space-propagated MSP-atlas (moving image) to the template-space, using the template MSP-atlas as the fixed image. This rigidly aligns an image containing only the midline of each sample with an image containing only the midline of the atlas, without any deformation, setting up an appropriate reference frame for measuring positional asymmetries like a protrusion asymmetry and overcoming the issue referenced by previous authors. We then applied this registration to similarly transform the full, sample-space annotations atlas for making positional measurements of each region. Our end result was the sample-propagated atlas being appropriately oriented within the reference/template space ("MSP-oriented full atlas") but only using information present at the tissue midline. This avoids any confounding asymmetry from the petalia in humans and any potential effect in other species.

In keeping with the surface-based approach from the prior studies, we converted the MSP-oriented full atlas to an outer-brain surface, from which we measured the mean protrusion of the frontal (anterior) and occipital (posterior) poles. We generated the outer surface by keeping only voxels where cortical brain tissue bordered universe voxels (empty space). In some non-human species, the most anterior or most posterior tissue was not the cortex (e.g. the cerebellum in chimpanzees or olfactory bulbs in mice), and so this tissue had to be removed from the sample's full regional atlas prior to generating the outer surface. Each surface was generated with the hemispheres digitally separated. We then selected the 1000 most anterior surface voxels in each hemisphere and the 1000 most posterior surface voxels for each hemisphere and calculated the mean anterior-posterior coordinate for both sides, for both poles. These values were our position measurements for the extent of the cortex in each hemisphere, and we calculated the protrusion as asymmetry of this position normalized against the length of the left hemisphere for that individual (linear distance between the left anterior and posterior pole). We used the equation:

$$petalia\ (frontal\ or\ occipital) =\ 100\% \times \frac{Right - Left}{Length\ of\ Left\ hemisphere\ for\ each\ individual}$$

We then transformed the raw data so that the typical petalia pattern in humans (right frontal more anterior and left occipital more posterior) was defined as a positive asymmetry. The 1000 voxels for each pole was approximately 2-4% of each hemisphere depending on the species, thus representing the most extreme elements of the tissue.

**Calculation of Positional Asymmetries.** For measuring positional asymmetries (mice only), we loaded each sample's MSP-aligned full atlas into python as a NumPy array and calculated the mean 3D centroid for each atlas region, within each sample. These were saved as .csv files for each sample which we loaded into R and transformed so that a right-anterior shift for a region was a positive asymmetry and a left-anterior shift was a negative asymmetry. In contrast to the petalia where we normalized the protrusion asymmetry by that *individual's* length of the left hemisphere, for simplicity, we normalized the positional asymmetries in each mouse sample by 173 voxels, the approximate length of the left hemisphere in our symmetrical CCFv3 atlas. White matter regions were dropped from the analysis as their diffuse structure limits their interpretation. We calculated the anterior-posterior positional asymmetry, for each region using the equation:

$$AP\ Positional\ asymmetry\ =\ 100\% \times \frac{Right - Left}{173\ voxels}.$$

We then performed similar statistics as with volume asymmetry – calculating cohort-specific mean positional asymmetry of each region, p-values, and performed cross-validation as described above.

**N4 Left-Right Reversal Experiment.** To assess whether the N4 bias correction algorithm functions in an intrinsically asymmetric manner (**Fig. S4E-F**), we performed the following procedure. 1) Load two copies of the same 3D image into memory. 2) To the first copy, apply the N4 algorithm. 3) To the second copy, flip the left-right axis, and then apply the N4 algorithm. 4) Reverse the left-right axis of the N4-corrected second copy, so that it matches that of the N4-corrected first copy (shown in **Fig. S4E**). 5) Subtract the N4-corrected second copy from the N4-corrected first copy (shown in **Fig. S4F**). This procedure should produce a Difference Image with all zeros if the N4 algorithm produced identical but flipped output images when given identical but flipped input images. However, the presence of non-zero values in the Difference Image indicates that this is not the case and is the reason that the correlation in **Fig. S4B** is lower than the expected R = -0.99.

**FreeSurfer analysis of human subjects.** Anatomical T1-weighted scans from 7 independent MRI datasets originating from 4 countries were used to compute asymmetry of cortical volume measures in humans. MRI images were processed with FreeSurfer[119] (v6.0.0), a fully automated procedure used to derive cortical morphometry surface maps for each MRI observation. One of the datasets was longitudinal and processed with FreeSurfer's longitudinal stream (datasets described in Roe et al., 2023[9]). Cortical volume maps for the left and right hemisphere were surface-smoothed using an 8 mm full-width, half-maximum Gaussian kernel and parcellated using the human connectome project multimodal atlas[120], which is well-suited for homotopic comparison. For cross-species comparisons and for analyzing the AP-pattern in volume (**Fig. 7, Fig. S9**), we calculated asymmetry as done for mice and NHPs:

$$asymmetry = 100\% \times \frac{Right - Left}{Left}.$$

For **Fig. S10**, initial volume asymmetry estimates were computed via the main effect of hemisphere in linear models controlling for age and sex, with a random subject intercept added where applicable. We then computed an asymmetry index (A.I.) as the above hemisphere effect divided by the intercept plus half the hemisphere effect. After correcting for covariates, this is equivalent to:

$$asymmetry = \frac{Left - Right}{(\frac{Left + Right}{2})}.$$

**Cross-Species Volume Asymmetry Comparisons.** To compare volume asymmetry between species, we used the Garin et al. 2022[82] multi-level, cross-species atlas. This atlas is a manually curated resource where atlas brain regions in 18 species are classified into a common organizational scheme. This was done at 4 different levels, starting most coarsely from "neocortex", "white matter", "cerebellum" and "brainstem" and going most finely to Brodmann segmentations, which are defined with cytoarchitecture. Since not every brain region of each species is matched with every region of every other species, especially at the finer levels, we calculated comparisons between each species using the regions that were shared in common between the 2 species in a given pair. This list of regions was thus different for each pair of species considered. For a given species, we used the multi-cohort asymmetry average calculated from all cohorts of that species in our study. In cases where multiple regions comprised a single larger structure, the mean volume of the comprising structure were summed for each hemisphere separately and then asymmetry was calculated from those summed volumes using the percent change formula used for **Fig. 7** and **Fig. S9**. For whole cortical lobes with aggregated subcortical structures, we used the level 2 annotations to create composite cortical lobes (frontal, parietal temporal, occipital), aggregated them into composite structures and calculated their asymmetry. We then repeated this process for subcortical structures (amygdala, claustrum, striatum, pallidum, hypothalamus, thalamus, colliculus, pons, medulla), which were available at level 3 (only using the subcortical structures which matched across the 2 species being compared). We then analyzed these combined sets of cortical and subcortical structures for **Fig. 7D**. For cytoarchitecturally-matched (Brodmann) regions

**(Fig. 7C)**, we used the same process but only using structures which matched across species at level 4.

**Exclusion of Chimpanzees from Volume Asymmetry Analysis.** Chimpanzees were excluded from region-wise volume asymmetry analysis because the high variability in the cortical folding pattern across individuals causes inaccurate mapping of cortical regions to the atlas when using the volumetric registration tools from our study (AntsPy, Elastix), even with the most flexible registration settings. As such, although our registrations were generally good, we could not fully align these secondary and tertiary cortical folds to the average template in a way that we felt would give a reasonably good mapping across species. Surface-based registrations, such as the one used for humans in Roe et al. 2023, are much preferred as they overcome the challenge of the varying cortical folds. However, such a pipeline that behaves symmetrically is not readily available for implementation in chimpanzees, especially one using the DaVi130 chimp atlas which is needed for cross-species comparisons. Hence, we proceeded without the chimpanzees in our cross-species comparison. We did however maintain the use of the Chimpanzees for measuring the anterior and posterior petalias, as this analysis does not depend on specific regional annotation but just on capturing the most anterior/posterior portions of the brain. We note that the volumetric registration tools work well with macaques and marmosets, as both species have cortical folding patterns which are the same across individuals.

**Establishing Left-right orientation in each Cohort.**

***Human cohorts.*** The Nifti header for all 4 cohorts was used to establish left-right orientation.

***Non-human cohorts.*** In contrast to humans, establishing the left-right (LR) orientation of the various mouse, marmoset, macaque, and chimpanzee imaging cohorts was complicated and was thus done with great care and in coordination with the original authors of the images. While Nifti headers are ideal for storing this information, they were only specified correctly on rare occasions, resulting in frequent mis-matches with the anatomy (see below for each MRI cohort). A description of how we established the LR orientation for each cohort follows below. In the STPT cohorts, file formats were non-medical imaging (.jpeg, .jp2, .tif) and thus no metadata information was available. Instead, we used information from tracer injections done by the labs who generated these images to define the left-right orientation.

*Mouse STPT Cohorts.* LR orientation of the mouse STPT cohorts was resolved through the use of tracer injections. For AMBCA, Oh et al. 2014[106] reported that a subset of animals from the AMBCA were injected in only the right hemisphere. We thus identified 50 of those animals within the cohorts and found that they all had clear and obvious injection sites on the same side, helping us to calibrate our analysis from here. For BICCN and Newmaster et al. 2020 cohorts, author Y.K. provided images of animals injected with tracers into a known hemisphere from both the Osten lab (BICCN; where he was previously a member and had knowledge) and his own current lab (Newmaster2020). The BICCN and Newmaster et al. 2020 cohorts were generated using different STPT machines at different institutions by different researchers. The BICCN images came from the Osten lab while the Newmaster et al. 2020 images came from author Y.K.'s lab.

*Pond and Qiu et al. 2018 cohorts.* Both cohorts were provided as .mnc files. Upon Mnc2Nii conversion (Minc-Toolkit V2), both had mis-specified nifti-header labels (and mnc header labels too). Neither cohort had the origin of real-world space at the anterior commissure. We thus used knowledge of the animal scanning orientation (POND: from author J.E., Qiu et al. 2018: from author

L.Q.) combined with knowledge of the MRI scanner's real-world coordinate system (provided by acknowledged individual J.S.) to identify the LR orientation.

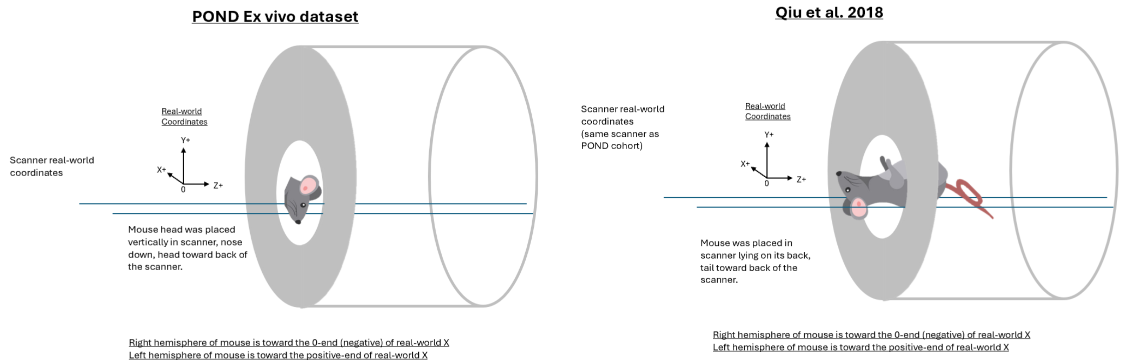


Of note, these 2 cohorts of mice were imaged by the same lab, using the same MRI scanner. However, most POND animals were raised and dissected for imaging in other labs and sent as fixed tissue to the Lerch lab, while the Qiu2018 animals were raised in-house by the Lerch lab.

*Guma et al. 2021 cohort.* This cohort was provided as .mnc files and upon Mnc2Nii conversion, was the only mouse cohort which showed proper alignments of the nifti header with the anterior-posterior and dorsal ventral anatomy. It also properly showed the anterior commissure set as the origin. Acknowledged individual G.D. confirmed that the .mnc files for the Guma et al. 2021 cohort had been specified properly according to the .mnc standard. Thus, properly converting to .nii (using Mnc2Nii) is expected to provide the correct anatomical header labels. We therefore placed confidence in these labels and used them for the LR orientation.

*Lopes et al. 2023 cohort.* This cohort was obtained as .nii files but the AP and SI axes were switched, although the anterior commissure was set as the origin. For this cohort, acknowledged individual E.K. directly provided the LR orientation description using a key that was initially provided by the first author A.S.

Overall, using these left-right assignments in mice, we found that asymmetries overwhelming went in the expected direction when compared between 2 given cohorts, when compared between the STPT and MRI average patterns, or compared during the cross-validation. Thus, while it is possible that we were wrong in our orientation for 1 cohort of animals, we view this possibility as unlikely, especially given that the STPT cohorts together show the same effect as the MRI cohorts.

*Macaque cohorts.* Acknowledged individual R.B. confirmed that the .nii header info was correct in the UW-madison cohorts. For the oxford macaque cohort, acknowledged individual M.R. confirmed that a right hemisphere marker was placed on each animal which was readily identified in the scans.

*Chimpanzee cohorts.* Animals had markers placed in known hemispheres which we readily identified in the images and which corresponded to information provided by author W.D.

*Marmoset cohorts.* The STPT cohort had tracers injected in only the left hemisphere which were readily distinguishable in the images. For the MRI cohort, acknowledged individual J.H. confirmed that the Nifi headers were correct for the left-right orientation.
